# Epigenetically bistable regions across neuron-specific genes govern neuron eligibility to a coding ensemble in the hippocampus

**DOI:** 10.1101/2019.12.13.874347

**Authors:** S. C. Odell, F. Taki, S. Klein, R. J. Chen, O. B. Levine, M.J. Skelly, A. Nabila, E. Brindley, J. Gal Toth, F. Dündar, C. K. Sheridan, R. N. Fetcho, A. Alonso, C. Liston, D. A. Landau, K. E Pleil, M. Toth

**Affiliations:** Department of Neuroscience, Weill Cornell Medicine, New York, NY, USA; Department of Pharmacology, Weill Cornell Medicine, New York, NY, USA; Physiology and Biophysics, Weill Cornell Medicine, New York, NY, USA; Applied Bioinformatics Core, Weill Cornell Medicine, New York, NY, USA; Department of Medicine, Weill Cornell Medicine, New York, NY, USA; New York Genome Center, New York, NY, USA

**Keywords:** Epigenetics, recruitment, neuronal excitability, dentate gyrus, DNA methylation, epialleles, neuronal heterogeneity

## Abstract

Episodic memories are stored in distributed neurons but how eligibility of individual neurons to coding ensembles is determined remains elusive. We identified thousands of predominantly bistable (CpG methylated or unmethylated) regions within neuronal gene bodies, established during the development of the mouse hippocampal dentate gyrus. Reducing DNA methylation and the proportion of the methylated epialleles at bistable regions compromised novel context-induced neuronal activation and spatial memory. Conversely, increasing methylation and the frequency of the methylated epialleles at bistable regions enhanced intrinsic excitability and spatial memory but impaired spatial working memory, indicating that the developmentally established methylated-unmethylated epiallelic balance at bistable regions is essential for proper neuronal excitability and hippocampal cognitive functions. Single-nucleus profiling revealed the enrichment of specific epialleles from a subset of bistable regions, primarily exonic, in encoding neurons. We propose a model in which epigenetically bistable regions create neuron heterogeneity, and specific constellations of exonic epialleles dictate, via modulating neuronal excitability, eligibility to a coding ensemble.

## Main

Most neurocomputational and experimental models predict a distributed neuronal coding scheme in which episodic memories are encoded by small and largely distinct ensembles of hippocampal neurons^1^. It is believed that sparsely distributed coding is an efficient way to produce high representational capacity with low interference. Sparse coding is particularly prominent in the dentate gyrus (DG) of the hippocampus because of the unusually low firing rates of DG granule cells (DGCs) and their higher number, relative to the number of their entorhinal cortex input and CA3 output neurons^2^. The first step in memory formation is the recruitment of a subpopulation of neurons to the coding ensemble. Neurotransmitters, paracrine signals, and hormones can “prime” neurons to exhibit a lower threshold for recruitment^3^. For example, synaptically released acetylcholine preferentially lowers the action potential threshold by inhibiting KV7/M current via increased axonal Ca^2+^, therefore enhancing intrinsic excitability and synaptic potential-spike coupling in DGCs^4^. However, the state of the neuron receiving these modulatory inputs is also essential^5^ because neurons with high intrinsic excitability are preferentially recruited during context exposure^6^. Excitability can be increased by a host of factors, from changes in the level and function of voltage-gated sodium and potassium channels to transcription factors such as activated CREB^7^. However, it is not known how these, or as of yet unidentified excitability-related molecules, are regulated to produce at any given time a small and continuously changing^8^ population of neurons eligible for coding.

Epigenetic processes, because of their cell-to-cell variability and dynamism, could underlie why a particular cell is recruited, while other neighboring cells are not, to a coding ensemble. Although the possible epigenetic basis of neuronal eligibility to a coding ensemble has not been studied, the ensuing process of memory formation has been associated with DNA methylation and histone modifications^9, 10^. Long term-potentiation (LTP) is dependent on de novo DNMT3A activity^11^ and histone actelyation^12^, and DNMTs are up-regulated in the hours following fear conditioning^13^. Epigenetic mechanisms are also vital to the maintenance and consolidation of memories, as administering DNMT inhibitors up to a month after memory formation results in failed memory recall^14^. Yet, to our knowledge, no study has investigated how epigenetic factors or epigenetic states may determine eligibility for recruitment, the first step of memory formation.

We found that CpGs in thousands of short regions in gene bodies exhibit high methylation/demethylation dynamics from middle/late embryonic stages in principal hippocampal neurons. This results in cells with different proportions of methylated and unmethylated epialleles and thus epigenetic mosaicism in adult neuronal populations. Bidirectional manipulation of the relative frequency of the methylated epialleles in vivo reduced and increased neuronal intrinsic excitability, novel context-induced neuronal activation, and hippocampal functions, linking methylation states and the combinations of these states, to neuronal eligibility to coding. Single-nucleus profiling of recently recruited neurons revealed enrichment for a set of epialleles that, via gene expression, may underlie eligibility for coding environmental inputs. Through these findings, our work implicates an epigenetic mechanisms dictating neuronal eligibility to recruitment, the first step of memory formation.

## Results

We found evidence for the developmental emergence of epigenetic heterogeneity in otherwise morphologically homogenous and genetically identical dorsal (d) DGCs in C57BL/6 male mice^15^. DGCs or their progenitors were microdissected from the granule cell layer of d (d) hippocampal slices at either embryonic day (E) 10.5, postnatal day (P) 6 or at 10-12 weeks of age, followed by enhanced RRBS methylation profiling of 1.5-2.5 million cytosines^16^ (Fig. 1a). Loss and gain of methylation during the transition of P6 young DGCs (yDGCs) to adult DGCs produced 170,198 differentially methylated (DM) CpG sites (q<0.01, RRBS at ≥10x coverage, average change 22.23%). In adult DGCs, these DM CpGs were by and large not fully methylated and unmethylated but rather were in the state of intermediate methylation (IM, i.e. were methylated across the entire 0% to 100% range, Fig. 1c, pink). Many of these sites were already IM at P6 (Fig. 1c**, blue**) indicating that the partial methylation/demethylation process began earlier. Indeed, comparing adult DGCs with cells from an earlier developmental stage, E10.5 hippocampal progenitors (HPs), still yielded DM sites (282,155 with an average change of 26.80%) which were IM in adult DGCs (**Fig.1b**, pink). These sites exhibited a more genome-typical bimodal methylation distribution in HPs (Fig. 1b, blue). IM in DGCs was unexpected because CpG sites in morphologically homogenous and genetically identical cells are typically uniformly methylated or unmethylated in all cells (i.e. the vast majority of CpG sites in the 0– 10% and 90–100% methylation range), resulting in a bimodal methylation distribution, as was the case at CpGs outside of the developmental IM CpGs (Fig. 1d,e).

**Figure 1:**
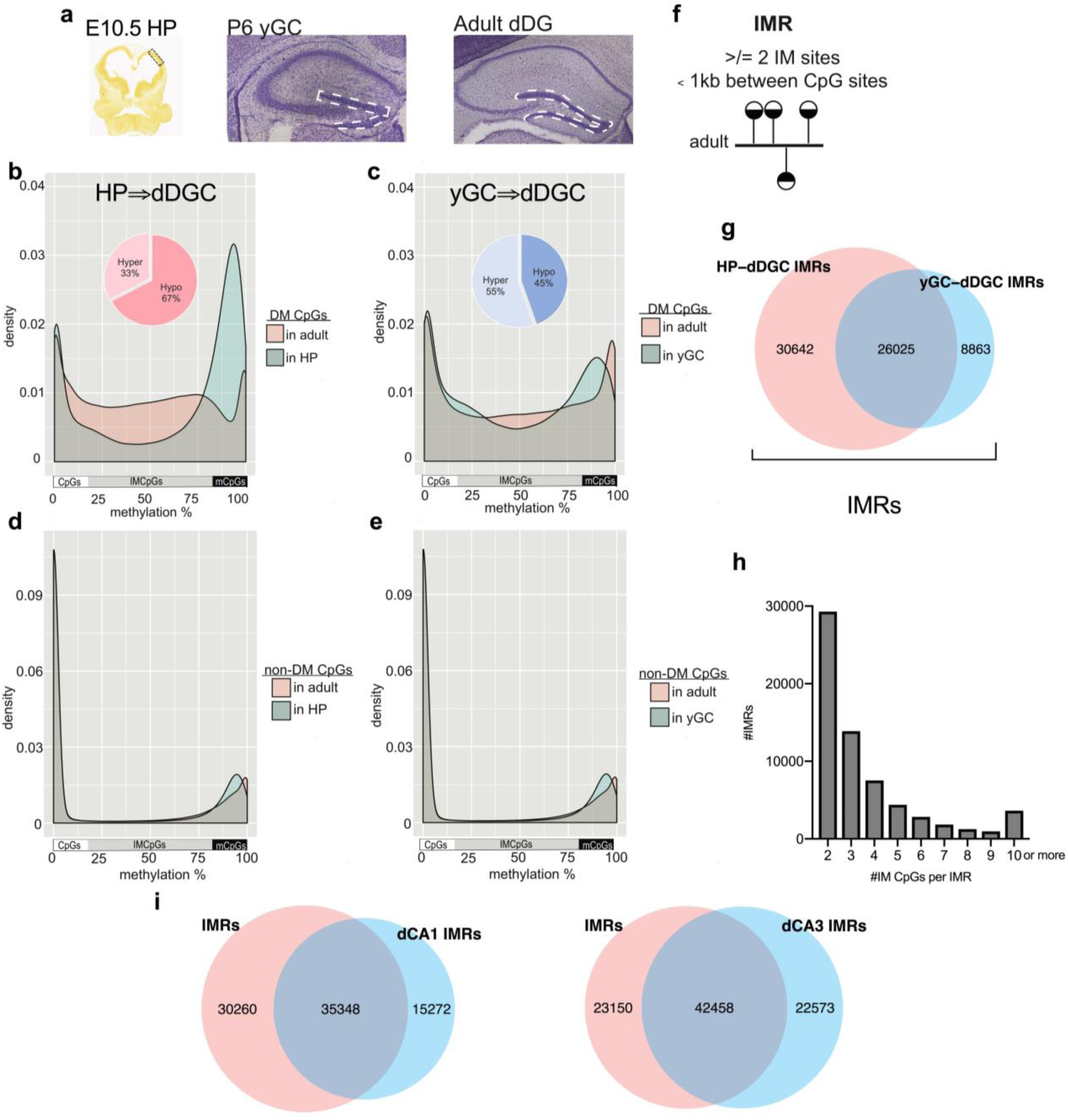
Emergence of genomic regions with intermediate methylation (IM) during hippocampal development. **a.** Cells used in DNA methylation profiling. HPs, yDGCs and adult DGCs were microdissected from E10.5, P6 and adult brains, respectively. **b,c.** Distribution of methylation levels at clustered (≥2) and differentially methylated (DM) CpGs between E10.5 HPs and adult DGCs (b) and between P6 yDGCs and DGCs (c). Sites initially (at E10.5) are mostly methylated and unmethylated indicated by their bimodal distribution, but by adulthood, they assume IM evidenced by a methylation pattern that covers the entire 0% to 100% range. While hypomethylation of methylated sites predominate during the transition of HPs to DGCs, methylation is slightly more dominant after P6. **d,e**. Non-DM sites have bimodal methylation distribution through development. **f**. IMR definition: regions with clustered IM CpGs in adult DGCs. **g**. IMRs encompass both HP-dDGC IMRs and yGC-dDGC IMRs. **h.** Distribution of IMRs according to the number of their IM CpGs. The largest group of IMRs contain 2 IM CpGs but ∼60% of IMRs have more than 2 IM CpGs. **i.** DM CpGs between 17.5 yCA and adult CA1/CA3 neurons are also IM in adult neurons (Supplementary Fig 1). They are clustered to CA1 and CA3 IMRs with an extensive overlap with IMRs identified in DGCs.

Around 80% of the IM CpGs, produced during the yDGC-to-DGC and HP-to-DGC transitions, clustered to IM regions (IMRs) of 2 or more IM CpGs, with an average size of 265.81 and 299.82 bps, respectively (Fig. 1f,g). The large overlap of yDGC-DGC IMRs with HP-DGC IMRs indicated selective DNA methylation/demethylation dynamics at the same CpGs throughout neuronal development, but the unique set of yDGC-DGC IMRs indicated additional methylation changes during DGC maturation (Fig. 1g). Because of the shared feature of IM at all these regions in adult DGCs, we henceforth used the combined set of IMRs (both HP-DGC and yDGC-DGC) in all further analyses (Fig. 1g, **Supplementary Table 1**, and **Supplementary Table 2** for links to custom tracks in Integrative Genomics Viewer). Overall, we identified 65,608 IMRs that on average contained 4.02 IM CpGs (Fig. 1h).

IMRs were not limited to DGCs in the hippocampus. Comparing the methylome of young CA (yCA) neurons (isolated from E17.5 embryos) with adult CA1 and CA3 cells microdissected from pyramidal layers yielded 50,620 and 60,031 regions exhibiting bimodal distribution in yCA cells but IM in adult CA1 and CA3 neurons, respectively (Supplementary Fig. 1a,b). DGC, CA1 and CA3 IMRs showed extensive overlaps consistent with their common origin and shared neurotransmitter phenotype (Fig. 1i).

The IM CpG profile of DGCs pooled from three males (displayed in Fig. 1b) was highly similar to that of six individual males (r^2^=0.89-0.90), which were also similar to each other (r^2^=0.93-0.94). This indicated that developmental IMRs are not the result of inter-individual variation in methylation, previously reported at transposon-derived sequences^17^ (Supplementary Figs. 2a,b). The high predictability of methylation levels at individual IMRs (Supplementary Figs. 2a) suggests that the IM pattern in adult DGCs arises stochastically during development. Importantly, IMRs are not related to genomic imprinting because they were not exclusively methylated around 50% (Fig. 1b,c pink) and did not map to classical imprinting control regions or non-canonical imprinted genes^18, 19^. IMRs are also different from so called partially methylated domains (PMDs), which are much larger regions spanning hundreds of kilobases and covering almost 40% of the genome of some cells^20^. Finally, IM in the adult granule cell layer was not due to differential methylation between developmentally and adult born DGCs as immature (<6 weeks old) neurons make up less than 10% of the total population size^21^, which is below the relative proportion of most methylated or unmethylated IMR CpGs in the DGC population. Furthermore, neurons in brain regions with no postnatal/adult neurogenesis (i.e. E17.5-adult CA1 and CA3) exhibited a similar IM pattern at a comparable number of IMRs (Supplementary Figs 1a,b). Overall, these experiments identified IMRs as a novel epigenetic feature in adult glutamatergic hippocampal neurons.

**Figure 2.**
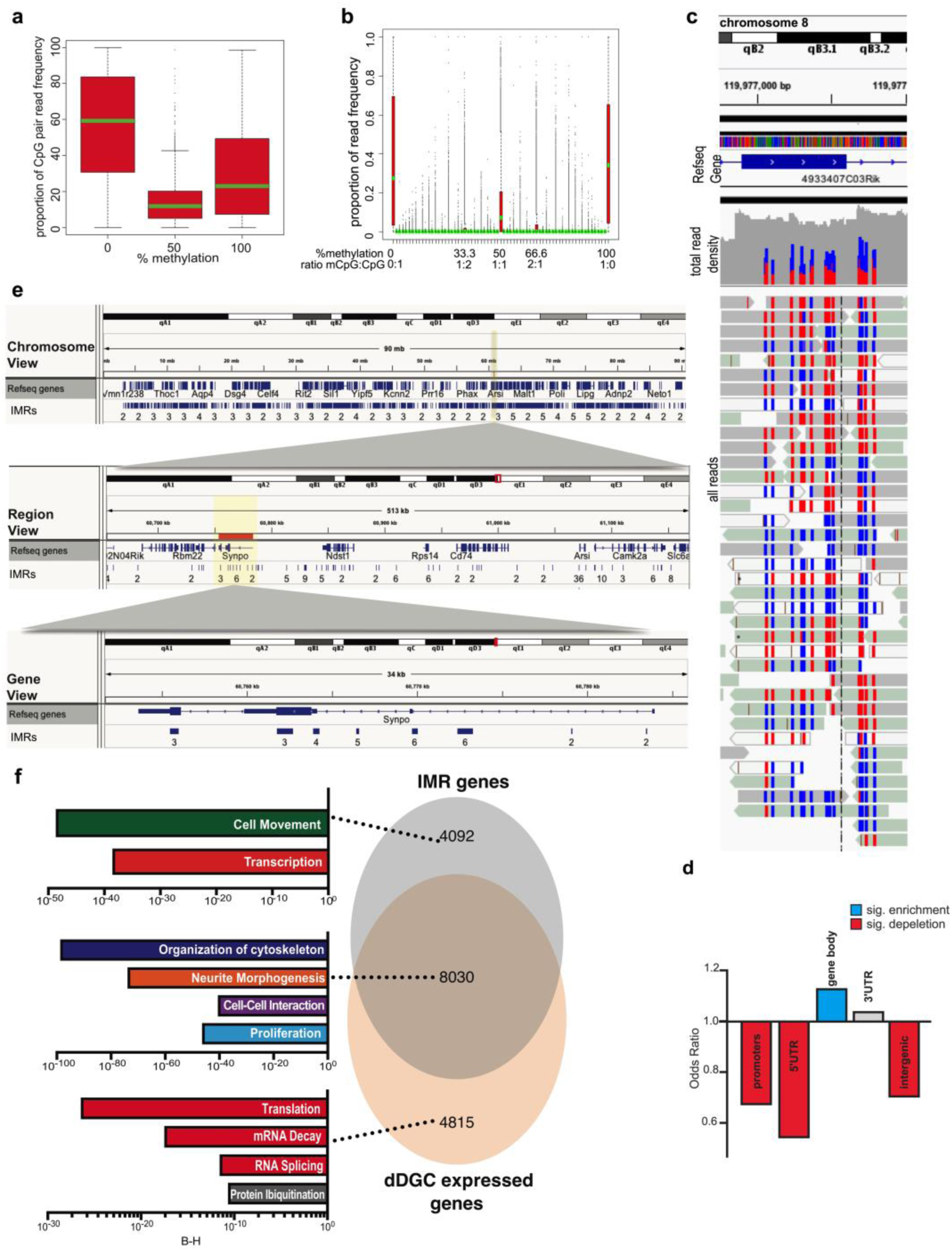
IMRs are largely bistable regions enriched in neuronal gene bodies. **a.** Two adjacent IM CpGs in adult DGCs tend to occupy the same methylation state. Means, SDs, and 5/95 percentiles are indicated. **b.** Neighboring IMR CpGs (∼4.37/IMR) in individual reads tend to be uniformly methylated or unmethylated indicating the epigenetically bistable nature of IMRs. **c**. Snapshot of individual reads of WGBS at an IMR shows a bistable methylation state, with most reads being either fully methylated or fully unmethylated at CpGs. RRBS produced a similar result. Blue: unmethylated CpGs, red: methylated CpGs. **d.** IMRs are depleted in promoters and intergenic regions but enriched in gene bodies. **e.** IGV browser view of *Synpo1* with custom IMR tracks overlayed at three different levels of magnification showing multiple IMRs spanning exons and introns. Numbers under regions indicate number of CpGs within IMR. **f.** DGC expressed IMR and non-IMR genes are enriched in different cellular functional categories (IPA, BH-corrected p values).

Analysis of individual sequence reads from DGCs (Fig. 2a) showed that two adjacent IMR CpGs more frequently occupy the same epigenetic state (positive correlation with Fisher’s test, adjusted p<0.05), than the two opposite states (Fig. 2b). Extending this analysis to multiple (>2) CpGs showed that neighboring CpGs (average of 4.37 IM CpGs/IMR) also tend to be uniformly methylated or unmethylated, but small fractions of IMR epialleles were epigenetically mixed, containing methylated (m)CpGs and CpGs, mostly in 1:2, 1:1 and 2:1 ratios (Fig. 2c). Methylation profiling also showed the emergence of IM cytosines in CA, CT, and CC dinucleotide contexts during DG maturation, but IM CpHs were not enriched at IMRs.

As some of the methylation signal in bisulfite sequencing (BS) can originate from 5-hydroxymethylated cytosines (5hmC), an intermediary between 5mC and C with potentially unique regulatory functions^22^, 5mC selective oxidative BS was performed (Supplementary Fig. 3a). However, 63.87% of IMRs were not hydroxymethylated, and those that were (36.13% of all IMRs) contained only 1.56 5hmCpG per IMR. Further, intermediately hydroxymethylated CpGs were not concentrated at IMRs (Supplementary Fig. 3b), therefore hydroxymethylation at IMRs likely reflects ongoing demethylation across the genome^23^ more than any specific role at IMRs. Finally, IMRs that contain hydroxymethylation retained their IM profile even after the removal of 5hmC sites (Supplementary Fig. 3c). Collectively, these data indicate that IMRs represent a unique set of short genomic sequences, characterized by heterogeneity in CpG methylation.

Next, we examined if developmental IMRs can be assigned to specific genomic features. IMRs had a relatively high, 6.59% CpG density, but were depleted in CpG islands (odds ratio 0.66, p<2.20E-16). Rather, IMRs were slightly enriched in CpG shelves (odds ratio 1.15, p=2.36E-9). Interestingly, IMRs were enriched in genic regions but depleted in intergenic areas (Fig. 2d). By dissecting genic regions we found that, although DNA methylation has a well-established role in transcription initiation, IMRs were depleted in promoters and 5’UTRs. Instead, IMRs were enriched in gene bodies, though exons, introns and last exon/3’ UTRs yielded no significant enrichment individually. The IGV view of representative IMRs within the gene body of *Synpo*, is displayed in Fig. 2e. This gene encodes the actin-associated protein synaptopodin which is thought to play a role in actin-based cell shape^24, 25^,. Methylation at gene bodies, though less well understood than at promoters, can regulate alternative splicing^26^ and 3D chromatin structure^27^.

IMR-containing genes represented ∼2/3^rd^ of all genes expressed in adult PROX1^+^ DGCs (Fig. 2f, Fig 5 a,b for FACS of PROX1 DGCs, **Supplementary Tables 3** and **4** for the list of IMR genes and expressed genes). Prox1 is a specific nuclear marker for DGCs^28^. IMR-containing expressed genes showed most significant enrichment in four main functions: I. *Cytoskeleton Organization*, II. *Neurite Morphogenesis*, III. *Cell-to-Cell Interactions/Contact* and IV. *Proliferation* (Fig. 2f, Ingenuity Pathway Analysis/IPA). The remaining ∼1/3^rd^ non-IMR expressed genes were associated with more universal cellular functions, such as *RNA Splicing*, *mRNA Decay*, *Translation* and *Protein Ubiquitination*. We also identified IMR genes that were not expressed/detected in our assay with functions in *Cell Movement* and *Transcription*. These genes may have been expressed earlier (e.g. cell movement) or expressed below the detection level in adult DGCs.

Given their DNA methylation based heterogeneity, association with gene bodies, and functional enrichments, we hypothesized that IMRs, via their combinatorial methylation states, “individualize” the structure and connectivity of DGCs, and that the resulting cellular heterogeneity is essential to support specific neuronal functions and behaviors. As reductionist methods such as gene targeting were impractical with the large number of IMRs, we sought to alter IMR methylation *en masse* while sparing methylation outside of IMRs. We previously observed that IMRs are more sensitive to changes in the methylation machinery than non-IMR sequences, presumably because of their bistability, as opposed to the monostability of non-IMR CpGs. Specifically, IMRs were sensitive to the gene dosage of *Dnmt3a,* encoding DNMT3A, the dominant *de novo* DNA methyltransferase expressed from mid/late fetal development^15^. Indeed, partial (∼75%^15^) and conditional deletion of *Dnmt3a* (cKO) in nestin-creERT2 mice^29^ by a single dose of tamoxifen (TAM) at E13.5 (Fig. 3a), a time point selected to interfere with the emergence of IMRs (Fig. 1b,c), resulted in the hypomethylation of a substantial number of CpG sites (18,825, q<0.01). These DM sites were IM in control cells (Fig. 3b, pink) and hypomethylated in cKO cells (Fig. 3b, blue). Importantly, CpGs that were not affected in cKO DGCs exhibited bimodal distribution and showed no appreciable leftward shift in their methylation distribution (Fig. 2c) indicating that hypomethylation in cKO cells was largely limited to IM CpGs. Notably, hypomethylated CpGs clustered to 4,194 differentially methylated regions (DMRs, ≥2 CpGs) that highly overlapped with the previously identified IMRs (85.96%, odds ratio=5.52, p<2.20E-16; Fig. 3d, Supplementary Fig. 4a, **Supplementary Table 2** for IGV track). Further, essentially all genes associated with cKO-DMRs (98.09%) were also IMR genes (Fig. 3e). However, in line with the partial nature of the cKO, not all IMR genes were affected, and only some of the IMRs within genes with multiple IMRs were hypomethylated (Fig. 2e). The partial and conditional deletion of *Dnmt3a* had no effect on the size of the DG and the extent of adult neurogenesis (assessed by the number of doublecortin^+^ neurons), indicating no apparent developmental effect of cKO on the number of adult DGCs (Supplementary Fig. 4b,c). Overall, these data indicate that IMRs can be hypomethylated, at least partially, by the conditional inactivation of *Dnmt3a*.

**Figure 3:**
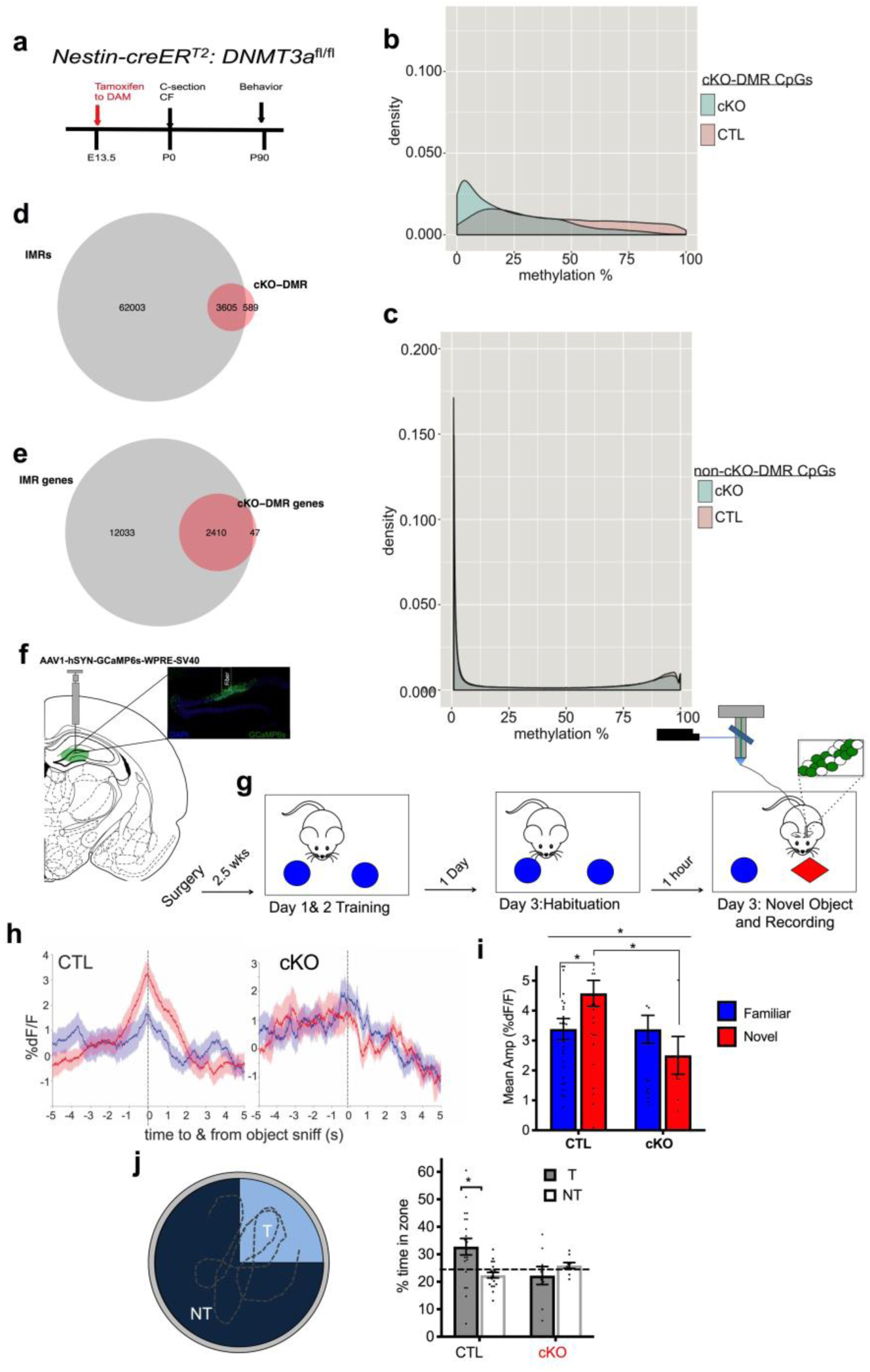
Emergence of IMRs during development is dependent on DNMT3A and is essential for proper neuronal activation and hippocampal dependent behavior. **a.** Timeline for generating *dnmt3a* cKO mice by TAM in *nestin-creER^T2^*-*dnmt3a^f/f^* mice. cKO (cre^+^/TAM), control (TAM). CF: crossfostering. **b.** cKO generates significant hypomethylation relative to control at select CpGs that can be clustered to cKO-DMRs (≥2 CpGs). Methylation of cKO-DMR CpGs in control DGCs is IM (pink), while is shifted towards 0% methylation in cKO DGCs. **c.** CpG sites with no significant methylation change in cKO neurons exhibit bimodal methylation distribution in both cKO and control neurons. **d.** Most cKO-DMRs are developmental IMRs. **e**. Essentially all cKO-DMR containing genes are IMR genes. **f.** Schematic of AAV1-hSYN-GCaMP6s injection and fiber implantation into dDG to record Ca^2+^ transients in the DG real time; inset: image of GCaMP6s expression and fiber implantation in dDG. **g.** Timeline of the novel object recognition test, with recording neuronal activity in the dDG by fiber photometry. **h.** Averaged traces of percent fluorescence changes in DG during interaction with novel object (red) or habituated object (blue), normalized to baseline (%ΔF/F) for cKO and control mice. Data are mean ± s.e.m. **i.** Quantification of average amplitude dF/F during object sniff of novel and familiar objects in control and cKO mice. The DG of control, but not cKO mice is activated in novel environment (Two-way ANOVA genotype x environment, F(1,88)=4.009, P=0.048; genotype, F(1,88)=4.073, P=0.047, N=5; 5). Significant difference between novel vs. familiar of control is also displayed: adjusted *p=0.032. Novel vs. familiar of cKO: p=0.320. Control vs. cKO in novel environment: adjusted *p=0.0134. Data are mean ± s.e.m. **j.** Percent of time spent in target (T) quadrant versus average of non-target (NT) quadrants of equal size in MWM during the probe trial on the fifth day in the absence of platform. cKO males fail to locate the platform, while control males recall the platform location (Two way ANOVA genotype x quadrant F1,54=7.133, P=0.0100; T vs. NT, Control and cKO males adjusted *p=0.0016 and p=0.405, respectively; N=cKO, 9; control, 20). Line represents chance.

**Figure 4:**
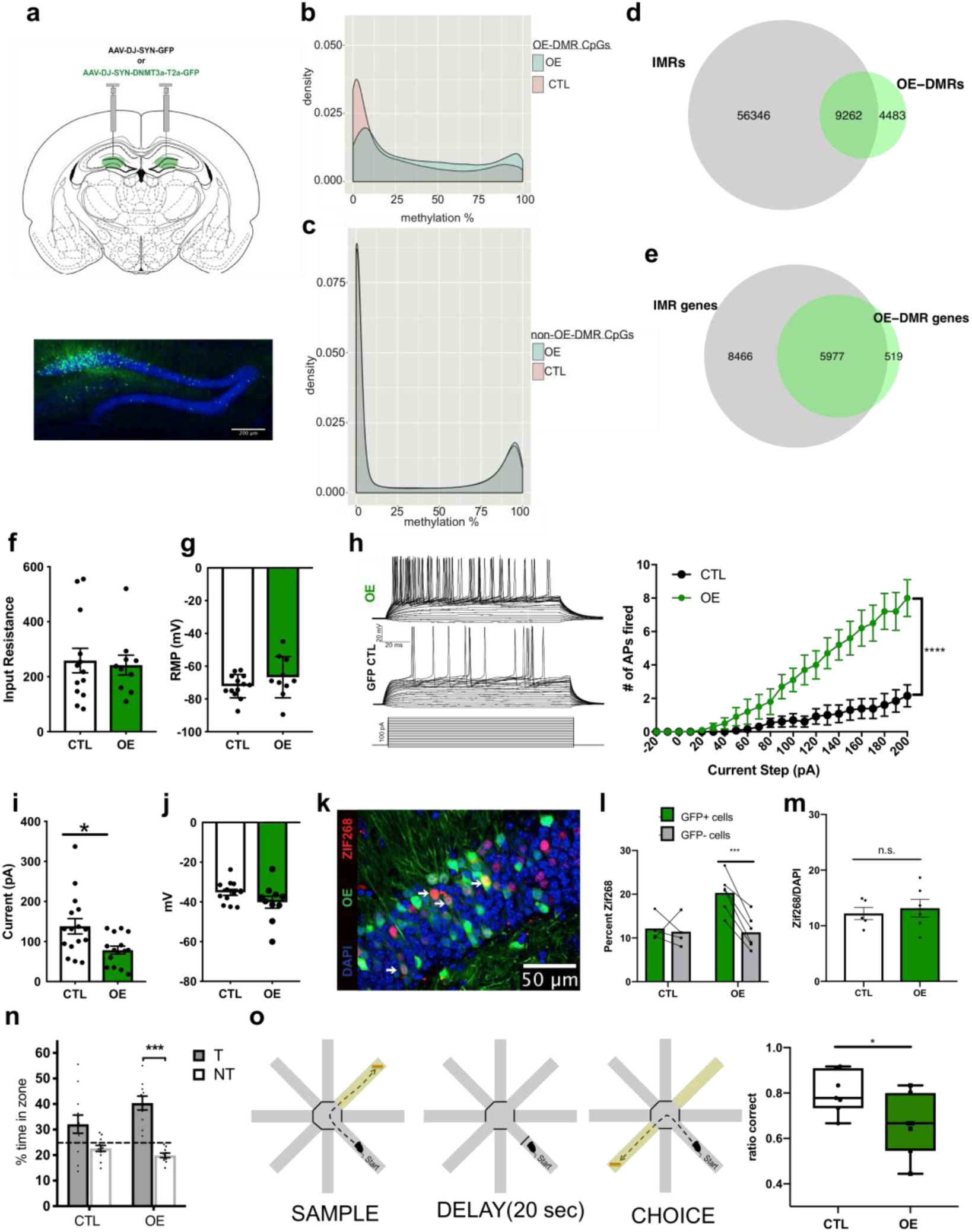
Overexpression of DNMT3A in DGCs increases methylation at a subset of IMRs, and enhances intrinsic excitability, hippocampal encoding and spatial memory, but impairs spatial working memory. **a.** Injection of DNMT3A overexpressing AAV-DJ-SYN-DNMT3a-T2a-GFP and control AAV-DJ-SYN-GFP into the dDG bilaterally. **b.** OE generates significant hypermethylation relative to control at select CpGs that can be clustered to OE-DMRs (≥2 CpGs). Methylation of OE-DMR CpGs in control DGCs is widely distributed from 0% to higher methylation. Methylation of these sites is shifted to the right in OE neurons. **c.** CpG sites with no significant methylation change in OE neurons exhibit bimodal methylation distribution in both OE and control neurons. **d.** OE DMRs show substantial overlap with developmental IMRs. **e**. Most of the OE-DMR containing genes are IMR genes. **f.** Similar input resistance of OE and control GFP^+^ neurons in DG slices (t=0.2783, p=0.7835; N=OE, 10 cells/6 animals; control, 13 cells/7 animals). **g.** Resting potential of OE and control GFP^+^ cells in DG slices (t=1.316, p=0.2023). Data are mean ± s.e.m. **h.** Slice electrophysiology recordings showing a significant increase in the number of evoked action potentials in OE as compared to control cells across a depolarizing current injection curve while cells are held at −70mV. Representative traces from an OE and a control cell are on the top and middle, current step protocol is on the bottom. Graphical representation of the data on the right (Two-way ANOVA genotype x pA F22,462=15.10, P<0.0001, genotype F1,21=14.85, ***P=0.0009, N=OE-D3A, 10 cells/6 animals, control, 15 cells/8 animals). **i.** Reduced rheobase in OE as compared to control cells (Welch-corrected t=2.754, *p=0.0114). **j.** Similar membrane potential at the occurrence of the first spike during current injection in OE and control neurons (t=1.573, p=0.1306). **k.** Representative immunohistochemistry image of dDG of mice after fifteen minutes of novel environment exploration. White arrows indicate OE and ZIF268 colabeled cells. **l.** OE GFP^+^ neurons were more likely to be recruited (ZIF268 positive) during novel environment exploration than their non-infected GFP^-^ neighbors, while control virus infected GFP^+^ cells were recruited at the same rate as non-infected GFP^-^ neighbors (Two way ANOVA GFP x experimental group F(1,8)=10.95, P=0.0107; GFP^+^ vs GFP^-^ in OE and control groups, adjusted ***p=0.0009 and 0.9182, respectively; N=OE, 6; control, 4, 2-4 sections per animal). **m.** Control and OE DGCs displayed similar levels of recruitment after novel environment exploration (t=0.426 p=0.6477, N=5, 6). **n.** OE mice spent significantly more time in target zone than non-target zones in the probe trial of MWM in the absence of the platform. While 4 days of training was not enough for control males to form memory (trend only), it was sufficient for OE males to recall the platform location (Two way ANOVA genotype x quadrant F(1,40)=5.150, P=0.0100; T vs NT in OE and control males, adjusted ***p=0.0003 and p=0.0708, respectively; N=OE, 10; control, 12). Also, control mice spent no more time in target quadrant than chance (One sample t-test: t(1)=2.716, p=0.2246). **o.** Performance of OE and control mice in the delayed nonmatching to place spatial memory task in the radial arm maze. Control mice outperformed OE mice (Welch-corrected t=2.331, *p=0.0407).

We next tested if the partial hypomethylation of IMRs is sufficient to interfere with coding in the hippocampus. We assessed neuronal activation, elicited by a novel object in a familiar environment, by monitoring population Ca^2+^ signal, a proxy for neuronal activity in the dDG by fiber photometry^30^ (Fig. 3f). Novelty is known to enhance extracellular glutamate levels in the hippocampus^31^ and to increase the firing rate and the frequency of Ca^++^ events of DGCs^31, 32^. Mice were exposed to two identical objects in three sample sessions in 3 days, followed on the last day by a test session with one familiar and one novel object. While control (TAM) mice displayed a robust increase in Ca^2+^ signal at the novel but not familiar object in the test session, no apparent increase in signal was detected at the novel (and familiar) object in cKO (cre+/TAM) mice, suggesting a failure of the dDG to detect/recognize novel object (Fig. 3g-h). However, novel object exploration and preference of cKO mice was not impaired and was similar to that of controls (Supplementary Fig. 4d), consistent with previous studies demonstrating spared novel object recognition after permanent hippocampal lesions due to compensatory changes in the perirhinal cortex^33, 34^. However, testing hippocampal functioning in the Morris water maze (MWM) revealed a deficit in spatial reference (long-term) memory. Specifically, in the probe trial of MWM, cKO mice, unlike their control littermates, explored all areas of the maze and had no preference for the quadrant that previously contained the platform (Fig. 3i). Overall distance travelled in the MWM was not different between cKO and control mice (847 and 963 cm, respectively; p=0.2784, t=1.106, df=27). We note that deletion of *Dnmt3a* postnatally (from ∼two weeks of age), after IMRs have been established, was reported to affect neither DNA methylation in the hippocampus nor spatial learning/memory in the MWM^35^. Thus, the spatial memory deficit of cKO mice can conceivably be explained by impaired developmental methylation, resulting in hypomethylation at IMRs and the subsequent defect in neuronal activation and coding of spatial information. Another hippocampal dependent behavior is contextual fear. Interestingly, the fear response of cKO mice was not different from that of controls during context only fear conditioning training and recall (in context A), and both were able to discriminate between training (context A) and new context (B) (Supplementary Fig. 4e), presumably because fear learning can be acquired via a non-hippocampal dependent strategy^36^. Collectively, these data indicate that expression of DNMT3A in developing neurons is essential for the emergence of IMRs and novelty-induced activation of DG neurons, with consequences on hippocampal dependent memory encoding.

Failure of DNMT3A deficient DGCs to respond to novel environment prompted us to test if the opposite, i.e. increasing DNA methylation by the overexpression of DNMT3A (OE) in adult dDG, increases the proportion of the methylated IMR epialleles and generates neurons with higher than average excitability. Again, we exploited the increased sensitivity of IMR CpGs to methylate them while avoiding the methylation of non-IMR CpGs (e.g. promoters and CpG islands) that are typically protected against de novo methylation^37^. Adult mice were injected with either DNMT3A expressing AAV-DJ-SYN-DNMT3a-GFP virus or control AAV-DJ-SYN-GFP virus bilaterally in the dDG, titrated to achieve a relatively sparse (10-30%) expression pattern (Fig. 4a). Twenty days later, both *Dnmt3a* and control virus-injected animals exhibited GFP expression in DGCs. Although limited to a fraction of cells, OE resulted in a population level hypermethylation at a large number of CpG sites (43,586, q<0.01). These CpGs, like IMR CpGs, displayed mostly IM in control cells (Fig. 4b, pink) and were relatively hypermethylated in OE neurons, indicated by a rightward shift in methylation (Fig. 4b, blue). CpGs that were not affected in OE cells exhibited bimodal distribution and showed no appreciable rightward shift in their methylation distribution (Fig. 4c) indicating that hypermethylation in OE cells was largely limited to IM CpGs. Notably, hypermethylated CpGs clustered to 13,745 DMRs (≥2 CpGs) that substantially overlapped with the previously identified IMRs (67.38%, odds ratio=2.85, p<2.20E-16; Fig. 4d, Supplementary Fig. 4a, **Supplementary Table 2** for IGV track). Further, the majority of genes associated with OE-DMRs (92.01%) were also IMR genes (Fig. 4e). Similar to hypomethylation by cKO, hypermethylation by OE was partial as not all IMR genes were affected, and only some of the IMRs within genes with multiple IMRs were hypermethylated (Fig. 2e). Overall, these data indicate that IMRs can be hypermethylated, at least partially, by the OE of *Dnmt3a*.

Next we tested if the partial hypermethylation of IMRs in DGCs is sufficient to increases their excitability. Since OE, as well as control cells were labeled by GFP, instead of population activity measures, it was possible for us to perform whole-cell patch-clamp electrophysiological recordings in brain slices. The input resistance of both OE GFP^+^ and control GFP^+^ neurons was similarly low (Fig. 4f), consistent with the 100-300 MΩ range reported for fully matured DGCs^38^. Although the resting potential (RMP) was not different (Fig. 4g), we found a significant increase in the number of evoked action potentials in OE as compared to control cells across increasing steps of depolarizing current injection, both at the common holding potential of −70mV (Fig. 4h) and RPM (not shown). In addition to this increase in current-injected firing, we found a decrease in the rheobase (minimum current required to elicit an action potential) in OE neurons using a current injection ramp (Fig. 4i), while the membrane potential at the occurrence of the first spike was not different between groups (Fig. 4j), together demonstrating that OE neurons are more excitable in response to depolarization than control neurons. Synaptic transmission, however, was unaltered in OE neurons, as the frequency and amplitude of spontaneous EPSCs and IPSCs were similar to control neurons (Supplementary Figs. 5a,b). These data demonstrate that overexpression of DNMT3A in DGCs enhances their propensity to generate action potentials in response to depolarizing current and presumably to excitatory synaptic inputs (i.e. increased intrinsic excitability).

As intrinsic excitability is thought to be a main factor driving neuron recruitment to an ensemble^39^, we next asked if OE cells have a higher than random chance to be included into a neuronal representation. OE mice (injected with AAV-DNMT3a-GFP) and controls (injected with AAV-GFP) in the dDG, were exposed to a novel environment for 15 min and 60 min later were sacrificed to visualize and calculate the fraction of DGCs positive for the immediate early gene (IEG) product ZIF268 in the infected (GFP^+^) and non-infected (GFP^-^) population of DGCs (Fig. 4k). OE-GFP^+^ neurons (i.e. neurons with increased intrinsic excitability) were more likely to be recruited (ZIF268^+^) during novel environment exploration than their non-infected GFP^-^ neighbors, while GFP^+^ and GFP^-^ neurons in control injected mice had equal probability for recruitment (Fig. 4l). However, there was no difference in the overall number of ZIF268^+^ DGCs relative to all DGCs (DAPI^+^) between OE and control mice suggesting that OE does not increase the recruited population size in the DG (Fig. 4m), presumably because strong GABAergic feedback inhibition limits the activated population to DGCs with the highest excitability^40^.

Recruitment of cells to an ensemble is the first step in learning/memory and thus increased neuronal excitability in the dDG of OE animals may influence hippocampal dependent tasks. Indeed, while 4 days of training in the MWM was not sufficient for the control mice to recall the platform, as they spent only slightly more time in the target than in non-target quadrants in the probe trial (trend only), OE mice had a strong and significant preference for the target quadrant (Fig. 4n). Overall distance travelled in the MWM did not differ between OE and control mice (1075 and 1024 cm, respectively; p=0.4574, t=0.7643, df=14). This demonstrated that OE animals formed a robust spatial reference memory, the exact opposite of the impaired memory of cKO animals in the MWM (Fig. 3i). Another form of hippocampal dependent memory is spatial working memory. This was tested in the delayed nonmatching to place paradigm in the 8 arm radial maze, employing opposite sample (S) and choice (C) arms with variable start arm, in a T maze-like configuration^41^. While control mice, as expected, strongly preferred the newly baited C arms to the already explored S arm, OE animals performed less well in this task (Fig. 4n) suggesting that OE of DNMT3A in a sparse neuronal population in the dDG reduces spatial working memory. Overall, our data show that either decreasing or increasing methylation incurs cost in cognitive functioning.

Since hypermethylation of IMRs in OE neurons was accompanied by increased neuronal excitability and hippocampal spatial memory, we predicted the methylated state of some IMRs to be enriched in neurons recruited to a coding ensemble. To test this notion, DGCs, activated by a 15 min exposure to a novel environment, were isolated 60 min later for single-nucleus genome-wide bisulfite sequencing (snRRBS). PROX^+^ DGC nuclei, expressing the IEG protein FOS, a proxy for neuronal activity, were sorted from PROX^+^/FOS^-^ cells from dissected dorsal hippocampi by FACS^28, 42^ (Fig. 5a). Novel environment, relative to home cage, increased the number of FOS^+^ neurons but, consistent with sparse encoding in the DG, FOS^+^ cells still represented only ∼2.3% of DGCs (Fig. 5b). The 264/288 FOS^+^ and 233/288 FOS^-^ sn-methylomes that passed quality control identified a total of 1,272,335 CpGs which were also present in the bulk DGC methylome (≥3 cells in both FOS^+^ and FOS^-^ populations, average 46.30 cells/CpG). Aggregate CpG methylation in FACsorted Prox1 cells, as determined by snRRBS, was highly concordant with methylation in microdissected DGCs by bulk RRBS (r^2^=0.8918). Direct comparison of methylation levels in FOS^+^ and FOS^-^ cells identified 14,881 DM CpG sites (1.16% of all detected CpGs, p<0.05, Bernard’s exact test, Supplementary Fig. 4a, **Supplementary Table 5**, **Supplementary Table 2** for link to custom tracks in IGV). Methylation at DM CpGs assigned individual FOS^+^ and FOS^-^ cells into two separate groups in t-distributed stochastic neighbor embedding (t-SNE) indicating that the FOS^+^ and FOS^-^ populations are relative homogeneous in their methylomes (Fig. 5c). Half of FOS-DM CpGs (49.4%) were preferentially in the methylated state (hypermethylated) in FOS^+^ cells vs FOS^-^ cells, while the other half (50.6%) were prefentially in the unmethylated form (hypomethylated) in FOS^+^ cells vs. FOS^-^ cells. Remarkably, in FOS^-^ cells, hypermethylated DM sites exhibited a wide range of IM distribution, similar to IMR CpGs in bulk DGCs (Fig. 5d, blue). These CpGs were in a higher IM range and even in the fully (23.1%, >90%) methylated state in the small population of FOS^+^ cells (Fig. 5d, orange). CpGs that are IM in FOS^-^ cells but are fully methylated (>90%) in FOS^+^ cells may represent methylated epialleles essential for, or at least compatible with, neuronal recruitment. Hypomethylated DM CpGs were also IM in FOS^-^ cells but contained a fraction of sites (27.2%) in the fully methylated state (Fig. 5e, blue), which were in the IM range in FOS^+^ cells (Fig. 5d, orange). The preferential exclusion of these fully methylated epialleles from FOS^+^ cells may indicate their incompatibility or interference with recruitment. CpGs whose methylation was not different between FOS^+^ and FOS^-^ cells, similar to non-IMR CpGs, exhibited a bimodal methylation distribution demonstrating that they were either fully methylated or unmethylated across all cells, whether FOS^+^ and FOS^-^ (Fig. 5f). These data indicate that, recruited/activated cells are different from the rest of DGCs at DM CpGs in the proportion of the methylated and unmethylated states. Some methylated CpGs were associated with FOS^+^ cells while others were dissociated from FOS^+^ cells.

**Figure 5:**
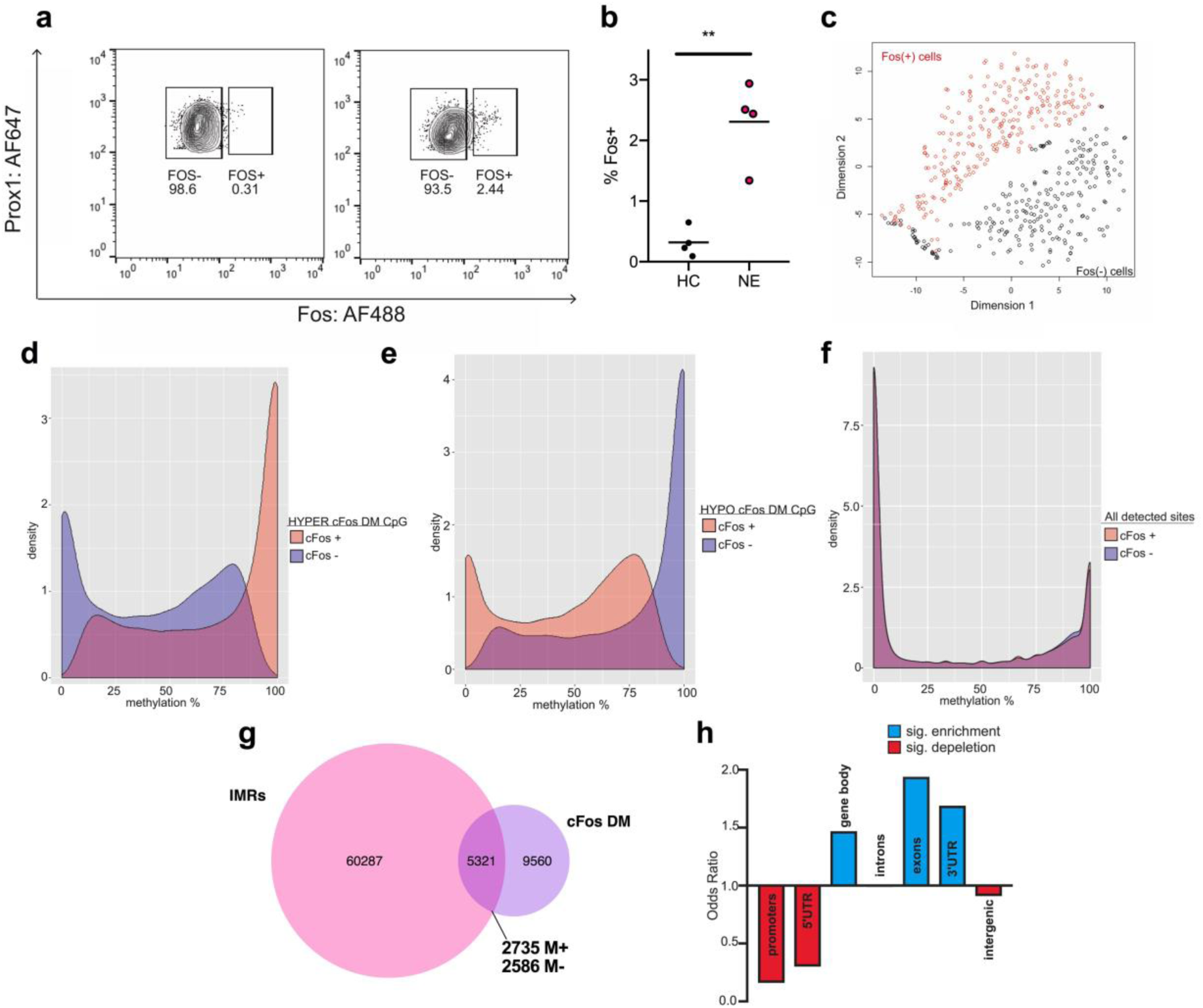
Sn-methylomes identifies DM CpGs between FOS^+^ and FOS^-^ cells that exhibit IM, map to developmental IMRs, and are enriched in gene bodies. **a.** Representative FACS plots showing the proportion of FOS positive nuclei in the population of PROX positive DGCs in home cage and after novel environment exposure. **b.** Proportion of FOS positive nuclei is increased by novel environment exposure. Four independent experiments (Welch-corrected t=5.504, **p=0.0066, 2-4 mice per group/experiment). **c.** T-SNE plot of FOS DM CpG sites shows the separation of FOS^+^ and FOS^-^ nuclei to two distinct populations. **d,e.** DM CpGs between FOS+ and FOS-cells, identified by snRRBS, exhibit IM across the entire 0% to 100% range. About half of FOS DM CpGs are hypermethylated in FOS^+^ (d) while the other half are hypomethylated in Fos^+^ (e). Number of DM sites are in “g”. **f.** Bimodal distribution of methylation levels at non-DM (1.2M) CpGs in FOS^-^ and FOS^+^ DGCs. **g.** Around 1/3^rd^ of DM CpGs between FOS^+^ and FOS^-^ cells mapped to IMRs. The methylation state of DM CpGs predicts the preference for the methylated (M+) or unmethylated (M-) IMR epiallele in the FOS^+^ population. **h**. IMRs associated with DM CpGs are depleted in promoters and intergenic regions but enriched in gene bodies, in particular exons and 3’UTR regions.

Since IM was a unique feature of both developmental IMR CpGs and DM CpGs, we next asked if they colocalize in the genome. We found that 36% of DM CpGs mapped to IMRs, a significant enrichment over chance (*p*=0.01291, Fig. 5g**, Supplementary Tables 1 and 5** for IMRs and FOS-DM CpGs, respectively). These IMRs were typically tagged with one DM CpG (1.21 DM CpG/IMR), likely because of the inherent sparsity of coverage by snRRBS and thus lower number of reads over CpGs relative to bulk RRBS (Supplementary Fig. 4a). Nonetheless, when two DM CpGs mapped to an IMR, they were typically uniformly methylated or unmethylated in FOS^+^, relative to FOS^-^ cells, much like neighboring CpGs in single IMR reads (Fig. 2a). This data indicated that the methylation state of DM CpGs may be used as a proxy to predict the preferred methylation state of their associated IMRs in FOS^+^ cells (Fig. 5g, M+ and M-). Further analysis showed that DM CpG-associated IMRs, similar to developmental IMRs, were depleted in promoters, 5’UTRs, and intergenic regions and were enriched in gene bodies (Fig. 5h). Interestingly, this group of IMRs was enriched in exons and 3’UTRs but not in introns suggesting possible sequence conservation.

Association of neuronal activation-related DM CpGs with developmental IMRs was much higher at gene level (80.2% of the 3,323 DM genes harbored IMRs), indicating the convergence of DM CpGs with a subset of IMRs on the same genes but not necessarily at the same location (Fig. 5i, top). Because of the IM state of most DM CpGs (Fig. 5d,e), this indicates the presence of additional IM areas within these genes that were not captured through developmental methylation changes between the three selected time points (Fig. 1a). Gene ontology by IPA showed the enrichment of all DM CpG genes (3,323), and those that also contained IMRs (2,665), in four functional categories (Fig. 6a with the top 10 functions and **Supplementary Table 6** for the list of genes), which were essentially the same as those we identified with IMR containing expressed genes (Fig. 2f): I. *Cytoskeleton Organization*, II. *Neurite Morphogenesis*, III. *Cell-to-Cell Interactions* (Cell-to-Cell Contact and Neuronal Activation/Transmission), and IV. Proliferation. Lack of enrichment of DM genes in RNA/protein expression/modification and metabolic processes, and in housekeeping genes in general, was notable suggesting enrichment in functions more specialized to neurons. Overall, two different approaches, one based on development and the other on neuronal activation identified IM CpGs/regions within neuronal gene bodies and implicate a role for these developmentally produced regions in neuronal recruitment/activation.

**Figure 6:**
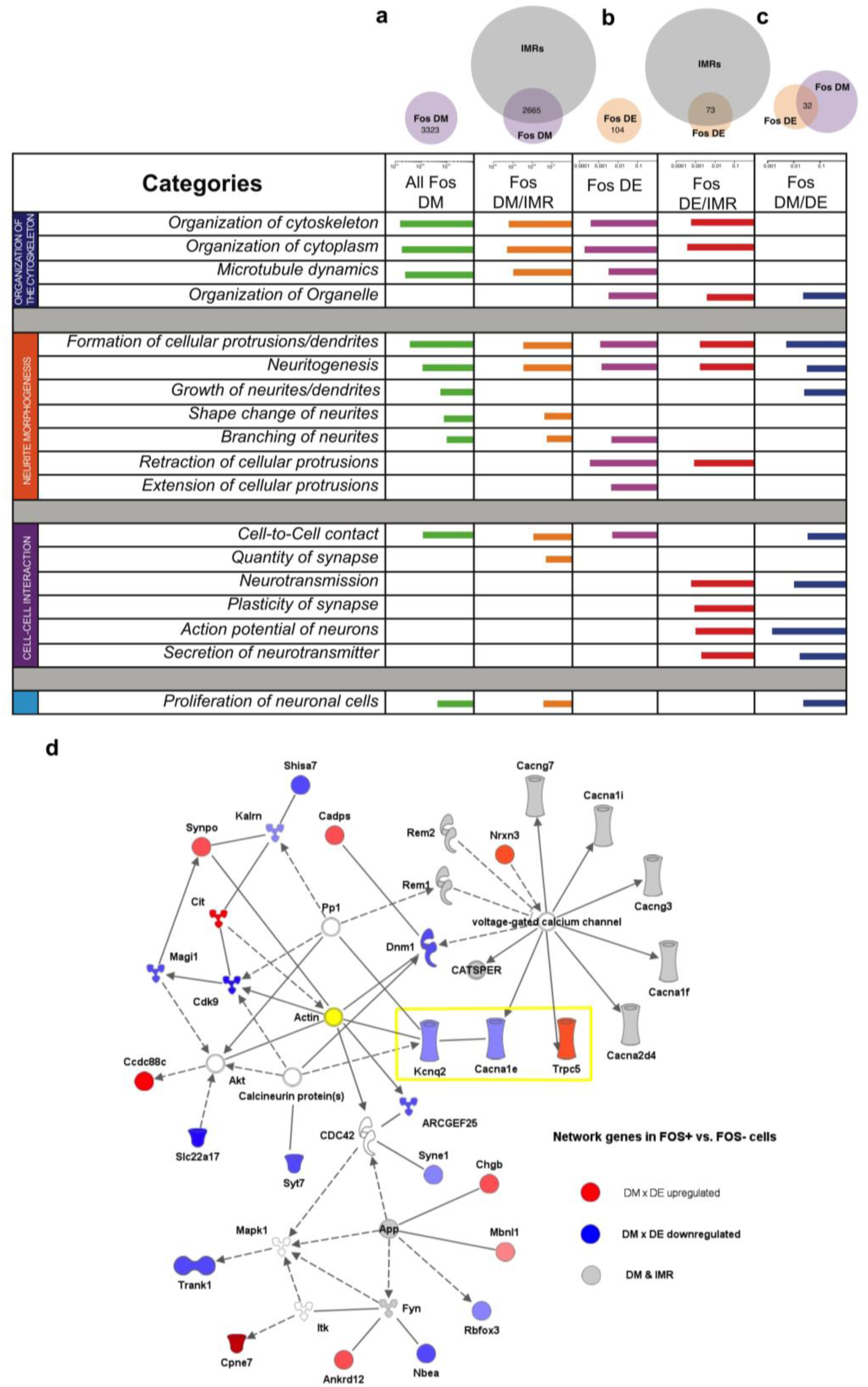
Sn-methylomes and transcriptomes of activated DGCs reveal common functional gene ontologies and networks. **a-c.** Ingenuity Molecular/cellular functions associated with DM (**a**), DE (**b**) and both DM and DE (**c**) genes between FOS+ and FOS-cells (IPA). Top ten functions are displayed based on BH corrected p values. Functions can be classified to four categories that are identical to those identified by expressed IMR genes in Fig. 2f. Three-quarters of DM x DE genes (c) are clustered to a gene network that includes several voltage gated channels and molecules interacting with actin/cytoskeleton suggesting regulation of intrinsic excitability and cell/neurite shape.

Next, we asked if genic DM between FOS^+^ and FOS^-^ cells is associated with differential expression (DE). We profiled 4273 PROX^+^/FOS^-^ and 721 PROX^+^/Fos^+^ DGC nuclei by single nucleus RNA-Seq (snRNA-Seq) using the 10x Genomics platform. Although 10x snRNA-Seq generates sparse gene expression profiles in individual cells (median # of genes 217/cell) we identified a total of 12,845 expressed and annotated genes (**Supplementary Table 4**). Of these, 104 were DE between FOS^+^ vs. FOS^-^ cells (adjusted p<0.05), 62.5% up- and 37.5% downregulated (**Supplementary Tables 7**). Most of the DE genes (72%) contained IMRs (odds ratio 1.35, Fig. 6b) and 32 genes (31%) were both DE and DM (odds ratio 83.35, p<2.2e-16 Fig. 6c**, Supplementary Tables 8**). Further, DE genes (Fig. 6b,c) were enriched in similar functional categories than DM genes (Fig. 6a), suggesting an epiallele-gene expression relationship. Specifically, enrichment of DE genes was seen in categories II. *Neurite Morphogenesis* and III. *Cell-to-Cell Interaction* (prominently *Neuronal Activity* and *Transmission*), while enrichment in functional categories I. *Cytoskeleton Organization* and IV. *Proliferation* was relatively diminished (top 10 functions displayed). Because individual genes had multiple IMRs and DM CpGs in gene bodies that each may regulate isoform-specific gene expression, it is challenging to establish a causative relationship between epialleles and gene expression at the single cell level. Moreover, given the large number of epialleles, their interaction may dictate the overall cellular phenotype and eligibility to recruitment. Therefore, we asked if genes which are both DE and DM (Fig. 6c) form an interactive network. A single IPA network clustered 75% of DE x DM genes, prominently represented by voltage gated potassium and calcium channels/subunits (*Kcnq2, Trpc5, Cacna1e*) that all can modulate neuronal excitability^43–45^ (Fig. 6d). The network also included genes encoding a calcium activated protein (*Cadps*), and a number of additional calcium channel genes that contained IMRs but were not DE (*Cacna1f, Cacna1l, Cacna2d4, Cacng3, Cacng7, Catsper 1,2,* and *4*). The prominence of calcium related genes in the network is in good agreement with the reported role of CREB, a cAMP/Ca^2+^activated protein, in neuronal recruitment^7, 46^ and is also consistent with the impaired Ca^2+^ signaling of cKO DGCs in which IMR methylation is perturbed (Fig. 3h,i).

Although genes encoding pre- and postsynaptic proteins (*Synpo, Nrx3, Kalrn, Syt7*) were in the network, we found no DE x DM genes specific for glutamate transmission, such as glutamate receptor subunits, suggesting the association of network genes with neuronal responsiveness to synaptic input, rather than with glutamatergic synaptic plasticity. This notion is consistent with the increased intrinsic excitability, but normal synaptic transmission of neurons in DNMT3A OE neurons, where the proportion of methylated IMR epialleles was increased (Fig. 4h). The network also contained genes related to small GTPase regulatory pathways which are known to regulate, via actin and cytoskeletal proteins, neurite/dendritic morphogenesis and plasticity^47^. ARCGEF25 and kalirin (*Kalrn*) are Rho guanine nucleotide exchange factors (GEFs), CIT is a Rho-interacting serine/threonine kinase, MAGI1 is a membrane associated guanylate kinase. REM1 and REM2 are two RRAD and GEM like GTPases. Synaptopodin (*Synpo*) is an actin binding protein and interacts with MAGl1 and kalirin. Overall, network genes defined interacting proteins regulating structural plasticity and neuronal excitability.

## Discussion

We identified thousands of small genomic regions, IMRs, that exhibit stochastic CpG methylation-based variability, predominantly bistability, within neuronal gene bodies. According to the theoretical model of Jenkinson et al.^48^, methylation bistability can be due to the low energy potentials of fully methylated and unmethylated epialleles, relative to alleles with a mix of methylated and unmethylated CpGs. Epiallelic bistability in hippocampal neurons is established by partial methylation and demethylation of unmethylated and methylated regions, respectively, during development. We recognized IMRs through developmental changes in methylation between 3 time points in the DG (E10.5, P6, and adult) and between 2 time points in the CA regions of the hippocampus (E17.5 and adult). However, additional IMRs are likely present in adult hippocampal neurons because of dynamic changes in methylation during development that were not be captured in our experiments. Indeed, snRRBS profiling of FOS^+^ and FOS^-^ cells, an approach that does not rely on developmental changes but rather on existing epigenetic heterogeneity and neuronal activation in the adult DGC population, revealed additional IM areas. Although these areas were mostly represented by single IM CpGs (due to the sparsity of coverage by snRRBS), many of them were within the boundaries of developmentally identified IMRs. Taken together, our data show the presence of IM and methylation bistability within neuronal gene bodies in adult hippocampal neurons by two independent approaches and connect developmentally produced IM sites/regions within gene bodies to neuronal recruitment.

Bistability in regional CpG methylation in bulk methylomes has been recognized in a wide range of human cells and tissue that was however primarily due to sequence-dependent allele specific methylation^49^. Sequence-independent (and non-parental) variability in methylation was reported in embryonic stem cells and progenitors suggesting their transient nature and role in early developmental processes^50, 51^. In contrast, the sequence-independent bistable methylation at IMRs described in this report is present in mature DGCs, implying a function in the adult hippocampus such as encoding of experiences. Indeed, our bulk methylome studies indicate that bistability at the developmentally established ratio between methylated and unmethylated epialleles is essential for neuronal recruitment for coding. First, reducing DNA methylation (by cKO of DNMT3A), and thus the proportion of methylated IMR epialleles, made neurons incapable of responding to novelty, and resulted in a spatial reference memory deficit in MWM. Beyond a deficit in encoding, cKO mice may have a deficiency in forming and recalling the memory of the platform location because learning and memory have been shown to depend on dynamic DNA methylation^9^. Second, increasing DNA methylation by the OE of DNMT3A, and thus the proportion of methylated IMR epialleles, caused hyperexcitability and impaired spatial working memory, but enhanced reference memory in the MWM. The artificially created hyperexcitable neuronal population in the DG of OE mice, although enhances long-term spatial memory, may be suboptimal to maintain network activity^52^ over the time scale of seconds to support spatial working memory.

Our single-nuclei based study showed that either the methylated or the unmethylated allele of a select group of IMRs is relatively enriched in activated neurons suggesting that methylation states at some IMRs may determine, or at least contribute to, neuronal eligibility to coding. Although our single-nucleus methylation data suggest that the preexisting methylation state of a subset of IMRs dictate neuron eligibility to recruitment, we cannot exclude the possibility that *de novo* methylation (and demethylation) during the short 60 min period between recruitment and immediate early gene expression plays a role. However, because IMR CpG sites that were preferentially methylated in FOS^+^ neurons were hypermethylated above chance (odds ratio=2.065, p=0.0018) in OE neurons (which are even more excitable), the preexisting methylation state of IMRs likely contribute to excitability and the neuron’s eligibility to recruitment. Overall, these data suggest a causal relationship between the methylation state of IM CpGs/regions within gene bodies and the propensity of the cell to be recruited to a coding ensemble.

Because IMRs were depleted in promoter regions, they may not directly regulate overall transcription. Instead, the exonic and 3’UTR enrichment of FOS-IMRs suggests a more complex DNA methylation dependent regulation that may include splicing, alternative 3’ length and consecutive changes in mRNA trafficking, subcellular compartmentalization and local translation^26, 53, 54^. Technical limitations do not currently allow mRNA isoform specific profiling at the single cell levels. Further, the presence of multiple DM CpGs/IMRs with occasionally opposing methylations states within a gene complicates establishing a direct relationships between epialleles and expression. Epigenetic editing^27^ could help to understand how individual epialleles influence spicing and isoform specific gene expression.

Although eligibility to recruitment to a neuronal ensemble probably relies on the state of multiple epialleles, some epialleles might be more central than others in generating the cellular phenotype. For example *Kncq2,* encoding a voltage gated K^+^ channel subunit in the interactive network (Fig. 6d), was downregulated in FOS^+^ neurons. Downregulation of this subunit alone could directly contribute to the higher excitability of activated cell because it is located in the axon initial segment^55^ and thus normally suppresses the initiation of action potential. Indeed, loss of KCNQ2 channels was reported to alter the intrinsic neuronal excitability and action potential properties of hippocampal and cortical neurons^43, 56–58^. Further, the majority of KCNQ2 epileptic encephalopathy variants are loss-of-function and dominant negative^59^ indicating that one copy of the variant is sufficient to alter the properties of the tetrameric potassium channel complex. *Trpc5,* an upregulated network gene in FOS^+^ cells that encodes a transient receptor potential ion channel regulating receptor- and store-operated Ca^2+^ entry, could be another central player in neuronal recruitment. TRPC is activated by G protein-coupled receptors, including muscarinic, adrenergic, PAR and serotonin receptors and thus its upregulation could enhance the effects of extrinsic modulatory signals on neuronal excitability^60^. Consistent with this notion, TRPC5 knockout mice exhibit significantly reduced seizure threshold and impaired long-term potentiation^44^. Finally, although the network gene *Cacna1e* encoding a voltage-dependent R-type Ca^2+^ channel subunit, was downregulated in FOS^+^ cells, the presence of additional Ca^2+^ channels and/or other molecules regulating Ca^2+^ homeostasis in the interactive network makes it difficult to predict the overall effect on excitability. Indeed, the presence of both up- and downregulated genes in the network is reminiscent of the dynamics of complex systems that can assume multiple stable states^61–64^.

Our model implicates IMRs, and specifically those defined by DM CpGs in activated neurons, in neuronal recruitment but not in the ensuing processes of learning and memory. Indeed, IMRs are not enriched in IEG genes, including *Ar*c and *Zif268*, that have been implicated in learning and memory^65, 66^. Neither other EIGs genes, such as *Myc* and *Jun*, or learning/memory associated genes, including *Fos*, *Bdnf,* and *Creb* have DM CpGs in activated neurons. Further, learning/memory associated increase in DNA methylation has previously been described in gene promoters, including the gene *Bdnf*^67^, while DM IMRs in FOS^+^ cells were found in exons and 3”UTRs. Therefore, our data indicate that the epigenetic mechanisms underlying neuron recruitment and learning are different. Recruitment may depend on stochastic epiallelic variability in a neuronal population to dictate intrinsic excitability and the probability of a neuron to be recruited to an ensemble. Learning/memory involves the activation of IEGs and epigenetic changes in recruited cells at specific genes to promote neuronal plasticity and Hebbian memory formation.

Although most of our studies were conducted with DGCs, we detected similar IMRs in other glutamatergic neurons in the hippocampus. This indicates that establishment of IMRs during neuronal development and their maintenance through adult life are a general phenomenon in principal neurons of the hippocampus. Therefore, we propose a model of neuronal recruitment in the hippocampus in which epigenetically bistable regions create neuron heterogeneity, and specific constellations of exonic epialleles dictate, via modulating gene expression and neuronal excitability, eligibility to a coding ensemble. Neuronal recruitment is the first and essential step in learning and memory, and our data in a mouse model demonstrate that disturbing its epigenetic regulation leads to a hippocampal encoding deficit.

## Acknowledgments

We would like to thank the following organizations for their support: The Epigenomics Core of Weill-Cornell Medicine, The Applied Bioinformatics Core of Weill Cornell Medicine, The University of Pennsylvania Vector Core, and the Stanford Vector Core. We would like to thank Paul Bergin for his guidance on behavior protocols and setup. We would also like to thank Dr. Thu Huynh for her guidance on novel object interaction behavior. Thank you to Chloe Lopez-Lee for your work on the hippocampal volume study. **Funding:** We are grateful for grant support **R01-MH103102 and R21-MH103715** from the NIH to M.T., Medical Scientist Training Program grant from the National Institute of General Medical Sciences of the National Institutes of Health under award number T32GM007739 to the Weill Cornell/Rockefeller/Sloan Kettering Tri-Institutional MD-PhD Program as well as by an NRSA from the NIMH under award number F30MH115622 to R.N.F

## Author contributions

Conceptualization, S.C.O., F.T., and M.T.; Methodology, S.C.O., F.T., R.J.C., M.J.S., A.A., C. L., D. A. L., K.E.P., and M.T.; Software, F.T., S.K., F.D., and R.N.F; Formal Analysis, S.C.O, F.T, M.J.S, R.N.F, K.E.P. and M.T.; Investigation, S.C.O., R.J.C, O.B.L, M.J.S, J.G.T, and C.K.S.; Writing, S.C.O. and M.T.; Supervision, A.A., D.A.L., K.E.P., C.L., M.T.

## METHODS

### Animals, intracranial virus injection and optical fiber implantation

Animal experiments were performed in accordance with the Weill Cornell Medical College Institutional Animal Care and Use Committee guidelines. All mice were group-housed two to five per cage, with a 12 h light/dark cycle and with lights on at 6:00 A.M. Food and water were available *ad libitum*. Wild-type C57BL/6 males were purchased from The Jackson Laboratory. Conditional *Dnmt3a* knock-out mice were generated as previously described^1^. C57BL/6 mice carrying floxed *Dnmt3a* alleles (*Dnmt3a^f/f^*^2^), provided by Riken BioResource Center, were crossed with mice heterozygous for the tamoxifen (TAM)-inducible nestin-cre-ERT2 transgene^3^, kindly provided by Luis Parada (University of Texas Southwestern Medical Center, Dallas, TX). To induce Cre-mediated *Dnmt3a* knockout at E13.5, pregnant cre-negative homozygous females were bred with cre-heterozygous males and were injected with 150-µl TAM solution (1mg), prepared by dissolving TAM (Sigma-Aldrich, St. Louis, MO) in ethanol and then in sunflower oil (Sigma-Aldrich, St. Louis, MO) (9:1) solution at 6.7 mg/ml. Because gestational TAM injection interferes with females’ maternal care behavior and labor, newborn pups were delivered via C-section on E20 and transferred into the nest of an early post-partum adoptive dam.

Intracranial fiber implantation into the dDG for fiber photometry. Male and female animals were anesthetized with 2% isoflurane and placed onto a stereotaxic frame to perform craniotomy. First, we delivered 275 nL viral vector AAV1.Syn.GCaMP6s at a rate of 75 nL per minute by a 10 µL Hamilton 700 series syringe with a 33 gauge blunt needle attached to a microsyringe pump and controller at the following coordinates: −1.94 mm anterior-posterior, −1.0 mm medial-lateral, and −1.85 mm dorsal-ventral. The needle remained at the injection site for 7 minutes post injection and was withdrawn slowly. A 400 µm diameter optical fiber was then implanted using the following coordinates: −1.94 mm anterior-posterior, −1.0 mm medial-lateral, and −1.83 mm dorsal-ventral. Optical fibers were secured with Metabond.

Intracranial virus injection to overexpress DNMT3A in the dDG. C57BL/6 males for virus mediated DNMT3A overexpression were purchased from Taconic Biosciences. Males were injected bilaterally, 500 nL/side with a rate of 50 nL/min into the dDG with either DNMT3A overexpressing virus, AAV-DJ-Syn-Dnmt3a-T2a-GFP (1.2E+07 IU./mL), or control virus, AAV-DJ-Syn-GFP (6.81E+08 IU/ml). Viruses were prepared by the Stanford Vector Core.

### Behavior

All tests were conducted using mice between 10 and 20 weeks of age. During all behavioral tests, the investigators who performed the tests were blind to the genotype and the treatment of the animals. Behavioral tests were conducted during the light on phase, between 1000 hours and 1600 hours. All mice were habituated to the behavior room for at least 1 h prior to testing. Mice were first tested for novel object recognition, followed by Morris water maze and radial arm maze tests, each spaced at least a 24hrs apart.

Novel Object Recognition. This behavioral test was performed in conjunction with fiber photometry to assess the activation of the dDG during familiar and novel object exploration. The test was conducted 2-2.5 weeks after the implantation of fibers and performed as previously described^4^. Different shaped and colored lego blocks (approximately 1 × 1 × 2 in) were used for the familiar and novel objects. Two days of training to familiar objects were performed in a 10-minute session, where male and female mice were exposed to the arena with the same two identical objects. On day 3, mice were again exposed to the same two identical objects in the arena for 10 min. One hr later, mice were reintroduced to the arena in which a novel object replaced one of the familiar objects on the non-preferred position (measured in previous session). Interaction with the objects for frequency, time, and duration was scored for 5 min from video recordings.

Morris Water Maze. Morris water maze was performed as previously described^5^. Mice were trained for four days with the platform in the NW quadrant of the maze. The probe trial was run 24h after the last training trial with the platform removed. Time in each zone was recorded for 60s.

Delayed nonmatching to place in Radial Arm Maze (RAM). Test was adapted from previously described^6^. Mice were food restricted one week prior to the start of testing and maintained at 85% body weight during testing. Mice were individually habituated to the maze for 10 minutes, where all arms were unblocked and the end of arms contained several food rewards (blue sprinkles). Then for 10 consecutive days, mice were tested in trials consisting of a sample and a choice phase. During the sample phase, all arms were blocked besides the arm with sample treat. The mouse was allowed to retrieve the food pellet in sample arm and was then brought back to the starting arm within 10 seconds after treat was retrieved. After 20 seconds in starting chamber, the choice phase was initiated, in which the choice arm, opposite to the sample arm and baited with sprinkle treat, was opened. Mice that entered the new choice (rewarded) arm, rather than the original sample (now unrewarded) arm were considered to have made correct choices. The sample arm was chosen at random. First day of data was considered training and not analyzed.

Novel Environment Exploration. Exposure to a novel environment was used to “mark”, by the expression of the early immediate protein FOS or zif268, the small population of “encoding” DG neurons for cellular and molecular studies. Mice were individually housed in dark boxes at least 2 hours prior to exposure to minimize background FOS/zif268 expression. Animals were placed one at a time into a large novel arena (24 x45 in) with many huts, tubes, and objects scattered throughout. Animals were allowed to freely explore for 15-minutes. Animals were then placed back in the dark for one hour to allow for maximum FOS or zif268 expression prior to transcardial perfusion or hippocampal dissection for FACs.

### Fiber Photometry

Fiber photometry was performed as previously described^7^ in the novel object recognition task as described above. To compare calcium signals across animals, fluorescence recordings were normalized to ΔF/F by taking the median signal across the recording period, subtracting it from each data point and then dividing by the median signal.

### Tissue preparation and immunostaining

Animals were sacrificed by transcardial perfusion with 4% paraformaldehyde (wt/vol), followed by one day post fix in 4% paraformaldehyde and at least 24 h in 30% sucrose cryoprotectant. Brains were flash frozen and were embedded in OCT followed by sectioning (40 µm) in a cryostat. For DCX cell counts, sections were incubated in 50% methanol for 30 min. Tissue was blocked in 1% BSA and incubated at 22–24 °C for 48 h with a goat polyclonal antibody to DCX (1:300, Santa Cruz, C18). Tissue was then incubated in secondary biotinylated rabbit antibody to goat antibody (1:250, Sigma, B-7024) for 2 h, followed by 1 h in ABC and DAB reagents for 2-10 min. Sections were counterstained with Nissl. DCX cell density in the dentate gyrus was measured using 4 sample sections (2 dorsal, 2 ventral) by using the StereoInvestigator software (MBF Bioscience). For granule cell layer (GCL) volume measurement, immunohistochemistry was performed by incubating the slices in DAPI (1:5000 Thermo Fischer D1306) for 10min in PBS. GCL volume was measured in 1:8 sections using the Cavalieri method by StereoInvestigator. For zif268 cell counts, tissue was blocked in 1% BSA and incubated at 4°C for 48 h with primary antibody, rabbit antibody to zif268 (1:500, Cell Signaling, 4153). Tissue was then incubated in secondary rhodamine antibody to rabbit antibody (1:500, Thermo Fischer, # 31670) for two h, followed by DAPI staining (1:5000) for 30 min. Dorsal sections were identified as between bregma −1.58 and −2.18 and were imaged at 40x on a Zeiss LSM 880 with each overexpressing virus image matched to a control virus section. DAPI, zif268^+^ and GFP^+^ cells were counted manually using the cell counter plug in FIJI. Cells were counted in each channel separately and then combined to identify GFP^+^/zif268^+^ colabeled cells. 2-4 sections per mouse were analyzed and counts were averaged per animal.

### Slice Preparation for recording

Mice were anesthetized with isoflurane and rapidly decapitated. Brains were removed and isolated in NMDG-HEPES artificial cerebral spinal fluid (aCSF) composed of (in mM): 93 NMDG, 2.5 KCl, 1.2 NaH2PO4, 30 NaHCO3, 20 HEPES, 25 Glucose, 5 sodium ascorbate, 2 thiourea, 3 sodium pyruvate, 10 MgSO4.7H2O, 0.5 CaCl2.2H2O (pH adjusted with HCl to 7.35, mOsm between 300 and 310), saturated with 95% O2 and 5% CO2. Transverse hippocampal slices (300 µm) were prepared using a Leica VT1200 vibratome (Leica Microsystems Inc., Buffalo Grove, IL), and incubated at 34 C for 11-12 min in NMDG-HEPES aCSF. Slices were transferred to a modified HEPES aCSF containing (in mM): 92 NaCl, 2.5 KCl, 1.2 NaH2PO4, 30 NaHCO3, 20 HEPES, 25 Glucose, 5 sodium ascorbate, 2 thiourea, 3 sodium pyruvate, 2 MgSO4.7H2O, 2 CaCl2.2H2O (Ph adjusted with NaOH to 7.35, mOsm between 300 and 310), and maintained at ambient temperature in oxygenated aCSF for at least forty-five mins before recordings began.

### Electrophysiological Recordings

Slices were transferred to a recording chamber and superfused with aerated aCSF containing (in mM): 124 NaCl, 2.5 KCl, 1.2 NaH2PO4, 24 NaHCO3, 5 HEPES, 12.5 Glucose, 2 MgSO4.7H2O, 2 CaCl2.2H2O at a rate of 2 ml/min using a perfusion pump (PeriStar Pro; World Precision Instruments, Sarasota, FL). For current clamp recordings of intrinsic membrane properties and current-injected firing, we used a potassium-gluconate–based intracellular recording solution containing (in mM): 135 K-Gluc, 5 NaCl, 2 MgCl2.6H2O, 10 HEPES, 0.6 EGTA, 4 Na2ATP, 0.4 Na2GTP, pH adjusted with KOH to 7.34, 280–290 mOsm. Spontaneous IPSCs and EPSCs (sIPSCs/sEPSCs) were recorded using a cesium-methanesulfonate-based internal solution containing (in mM): 135 Cs-Meth,10 KCL,10 HEPES, 1 MgCl26H2O, 0.2 EGTA, 4 MgATP, 0.3 Na2GTP, 20 phosphocreatine, pH adjusted with CsOH to 7.34, 280-290 mOsm. Recording electrodes were prepared from filamented borosilicate glass capillary tubes using a horizontal micropipette puller (P-1000; Sutter Instruments, Novato, CA). Whole-cell patch-clamp recordings were collected from neurons voltage-clamped at −55 mV for sEPSCs and +10 mV for sIPSCs; current clamp experiments were conducted at the cell’s resting potential and at −70 mV. A maximum of two cells per experimental condition were recorded from each animal, and all cells included in these analyses maintained a stable access resistance of 5–20 MΩ. Whole-cell currents were acquired using a MultiClamp 700B amplifier, digitized (Axon Digidata1550B, Axon Instruments, Union City, CA), and analyzed online and offline using an IBM-compatible personal computer and pClamp 10.3 software (Axon Instruments).

### Flow cytometry

Nuclei were isolated as previously described^8^. Briefly, the whole brain was sectioned coronally using a mouse brain slicer matrix to isolate the dorsal hippocampus. The dorsal hippocampus was dissected out and placed in a nuclei isolation medium (sucrose 0.25 M, KCl 25 mM, MgCl_2_ 5 mM, TrisCl 10 mM, dithiothreitol 100 mM, 0.1% Triton, protease inhibitors). Tissue was mechanically homogenized, allowing for separation of nuclei from cells. Samples were washed, resuspended in nuclei storage buffer (0.167 M sucrose, MgCl_2_ 5 mM, and TrisCl 10 mM, dithiothreitol 100 mM, protease inhibitors) and filtered through 70 µM then 30 µM filters. Solutions and samples were kept cold throughout the protocol. Dissociated nuclei were then stained by 30 min incubation in anti-Prox1 Alexa Flour 647 (NBP-1-30045AF647, NovusBio, 1:1000) and anti-c-FOS Alexa Flour 488 (NBP2-50037AF488, NovusBio, 1:1000). Samples were gated and collected using a BD Influx sorter followed by analysis using the FlowJo software (Tree Star). Nuclei were gated first using forward and side scatter pulse area parameters (FSC-A and SSC-A) excluding debris, followed by exclusion of aggregates using pulse width (FSC-W and Trigger Pulse Width). The FOS^+^ population was gated using home cage control populations based on PROX1 and FOS fluorescence.

### DNA Collection for bulk methylation sequencing

Brains from adult WT, transgenic, and virus injected mice were collected, quick frozen on dry ice and sectioned (200 µM) in cryostat. The dentate gyrus, CA1 and CA3 were microdissected from the sections. DNA was isolated using the QIAamp DNA Micro Kit according to manufacturer instructions (Qiagen).

### Reduced Representation Bisulfite Sequencing (RRBS)

In most experiments, we used RRBS^9^ because of its high efficiency in sequencing CpGs and its higher coverage of sequenced reads relative to that of whole genome bisulfite sequencing (WGBS) (160X vs. 30x) that allowed us to more accurately determine the extent of developmental methylation changes. Preparation for RRBS libraries, sequencing, and adapter trimming were done by the Epigenomics Core at Weill Cornell Medicine. Single end 50 bp RRBS sequencing was performed as described, using Illumina HiSeq2000 and 2500 machines according to the manufacturer’s instructions. UCSC mm9 was prepared and indexed followed by alignment and methylation calling using Bismark (v17) (Akalin et al., 2012b). Differential methylation and statistical analysis were performed using the MethylKit package in Differential methylation and statistical analysis were performed using the MethylKit package^10^ in R at default setting. Differentially methylated sites were defined as sites where the sliding linear model (SLIM)-corrected p-values (q) were ≤0.01. Differentially methylated regions were defined as regions containing at least two differentially methylated CpG sites with <1 kb distance between sites, and were referred to as an intermediate methylated region (IMR). Bed files for CpG islands were prepared using a publically available pipeline (https://www.r-bloggers.com/cpg-island-shelves/). Genomic coordinates for exons, introns, 5’UTR, 3’UTR were downloaded from the University of California Santa Cruz (UCSC) genome browser based on mm9. Promoters were defined as regions ±500 bp from the transcription start site (TSS). To determine if IMRs are enriched in a genomic feature, we computed the Odds Ratio as: [(the number of IMR overlapping feature)/(the number of IMR not overlapping feature)] / [(the number of potential IMR overlapping feature)/(the number of potential IMR not overlapping feature)]. Odds Ratio >1 reflects enrichment of the IMR to feature, while Odds Ratio <1 reflects depletion. *To get the potential DMRs in the genome outside of the identified DMRs, we clustered all detected non DMR CpG sites DMRs (4 sites, neighbors ≤ 1kb) (PMID: 30643296).* Statistically significant enrichments were computed using Chi-squared test in R. Density plots were done using ggplot function (ggplot2, R).

### Whole Genome Bisulfite Sequencing (WGBS)

Single-end 50 bp WGBS sequencing was performed using Illumina HiSeq2000 and 2500 machines according to the manufacturer’s instructions. The pipeline published by Lister and coworkers^11^ was used for alignment, methylation calls, and differential methylation.

### Reduce Representation Oxidative Bisulfite Sequencing (RRoxBS)

To identify cytosines with hydroxymethylation (5hmC), RRoxBS, in combination with RRBS, was performed as previously described^12^ with modifications for reduced representation sequencing. Library preparation, sequencing, and adapter trimming were done by the Epigenomics Core at Weill Cornell Medicine as follows: Briefly, genomic DNA (100 ng/sample) was digested with 100U of MspI (New England Biolabs, Ipswich, MA), and cleaned up using QIAquick PCR purification columns (QIAGEN, Hilden, Germany). The digested DNA was spiked with 0.01% control DNA duplexes, provided by Cambridge Epigenetix (Cambridge, UK), containing C, 5-fC, 5-mC and 5-hmC bases at known positions. The control sequences were used to give a quantitative assessment of the efficiency of oxidation and bisulfite conversion. The rest of the library preparation was carried out using the Ovation Ultralow Methyl-Seq DR Multiplex with TrueMethyl oxBS kit and workflow (Tecan, Redwood, CA). The libraries were split into two aliquots after adapter ligation: one aliquot was oxidized and bisulfite converted (oxBS), the other subjected to a mock oxidation before bisulfite conversion (BS). 2ul of the resulting material was assessed by qPCR to determine the optimal number of PCR amplification cycles. After amplification, the libraries were normalized and pooled according to the desired plexity, clustered at 6.5pM on single read flow cells and sequenced for 50 cycles on an Illumina HiSeq 2500. Illumina’s CASAVA 1.8.2 software was used to perform image capture, base calling and demultiplexing. Analysis of bisulfite treated sequence reads was carried out as described^13^, using CUTADAPT instead of FLEXBAR to trim the reads. Alignment to the mm10 genome and methylation calls were done using Bismark (v17) using the default options. Global CpG report was then used to compute percent CpG methylation and total coverage with base-pair resolution. To identify the level of 5hmC at a CpG site, we considered only the sites that are detected in the BS and oxBS experiments with delta methylation (BS – oxBS) being > 0. The BS experiment does not discriminate between 5mC and 5hmC, while the oxBS experiment detects 5mC. Therefore, the 5hmC methylation levels were inferred by subtracting oxBS methylation proportion from BS methylation proportion. Proportion test (prop.test, R) was then used to compute statistically hydroxyl-methylated (5hmC) cytosines. Therefore, CpGs with statistically significant differences between BS and oxBS experiments (p < 0.05, methylation proportion (BS – oxBS > 0)) were considered significantly hydroxy-methylated (PMID: 28428825).

### Multiplexed single-nucleus RRBS (snRRBS)

Single nucleus RRBS were performed by first sorting PROX^+^/FOS^+^ and /FOS^-^ neuronal nuclei in 96-well plates in 3 µL of 0.1× CutSmart buffer (New England Biolabs) per well as described in the Flow Cytometry section of the Materials and Methods. Plates were stored at −80 °C. Processing of samples were as described for Multiplexed single-cell RRBS^14^ by the Epigenomic Core Facility of Weill Cornell Medicine. Briefly, nuclei were lysed, DNA was cut with Msp1 (Fermentas), and A-tailed DNA fragments were ligated with custom methylated adapters containing inline cell barcodes. Then, adapter-ligated DNA from 24 nuclei were pooled together based on compatible cell barcodes, yielding 4 pools of 24 individual nuclei per 96-well plate. Each 24-nuclei pool was subjected to two cycles of bisulfite conversion (EpiTect Fast Bisulfite Kit, Qiagen) according to manufacturer’s instructions. Converted DNA was then amplified using primers containing an Illumina i7 index. A different i7 index was used for every 24-nuclei pool, allowing multiplexing of 96 nuclei for paired-end sequencing on one Illumina HiSeq 2500 lane.

Each pool of 96 nuclei was first demultiplexed by i7 barcodes, and then each pool of 24 nuclei was further demultiplexed by unique cell barcodes. Processing of the reads was adapted from (Gaiti, et al). Briefly, reads with at least 80% match for the template adapters were assigned to a given cell. Adapters were trimmed from the raw sequence reads. After adaptor removal, the first 5bp were trimmed from the 5’-end of the R1 and R2 fastq files. Reads were then aligned to the mm9 mouse genome using Bismark34 (v.0.14.5; parameters: -multicore 4 -X 1000 -un -ambiguous) using bowtie2-2.2.8 aligner35. Methylation states at each CpG were deduced using Bismark methylation extractor (-bedgraph -comprehensive). A total of 264/288 FOS^+^ and 233/288 FOS^-^ cells passed the filtering criteria that included minimal coverage of 50,000 unique CpGs per cell, bisulfite conversion rates ≥ 99%, and number of reads below the 99^th^ percentile of coverage in the cell. Collapsed methylation signals specific for the cell and site were saved as a matrix and inputted into R for statistical analysis and for data visualization. A total of 1,272,335 CpGs, which were also present in the bulk DGC methylome (≥3 cells in both FOS^+^ and FOS^-^ populations), were detected across FOS^+^ and FOS^-^ cells.. Site differential methylation (DM) between FOS^-^ and FOS^+^ cells was done using Barnard’s exact test (p<0.05) (Barnard package, R) for all sites. Site coordinates were overlapped with other features using intersectBed tool. The methylation profiles across all FOS^+^ and FOS^-^ cells was done using Rtsne package in R. For density plots, the average methylation per site was computed by dividing the cell counts for 100% methylation by the total number of cells within which a site was detected. The data was then used for density plots using ggplot2 in R. Odds Ratio for enrichment above chance was computed by (A=the number of sites overlapping feature/B=total number of sites)/(C=same number of random sites overlapping feature/D=total number of sites). This process was repeated 10 times. The average Odds Ratio was then computed and used for Chi-squared analysis in R.

### Single nuclei RNA-Seq on 10x Genomics platform

Single nucleus RNA-seq were performed by first sorting PROX^+^/FOS^+^ and /FOS^-^ neuronal nuclei in separate tubes of PBS as described in the Flow Cytometry section of the Materials and Methods. Cell suspension were kept on ice and transferred to the Weill Cornell Epigenomics Core Facility for RNA-Seq. Standard 10x Genomics Chromium 3′ libraries (v3) were prepared according to the manufacturer’s recommendations. The FOS^+^ and FOS^-^ 10x libraries were sequenced together on a HiSeq 2500 (Illumina) machine. 10x data were processed using CellRanger 3.1.0 with default parameters. Reads were aligned to the mouse reference genome mm10.

The CellRanger aggr pipeline was used to aggregate FOS^+^ and FOS^-^ libraries into a single gene expression matrix. This matrix was loaded into R using the Seurat package (version 3.1.1) for downstream analysis. Default parameters were used unless specified. Nuclei were excluded from analysis if: 1) fewer than 200 genes were expressed 2) greater than 2500 genes were expressed or 3) more than 20% of UMIs were mapped to mitochondrial genes. Genes not expressed in at least three nuclei were also filtered out. Gene expression was normalized to the total expression per nucleus, scaled by 10,000 and log-transformed. Mitochondrial genes were regressed from each nucleus and gene expression was scaled. Differential expression between FOS+ and FOS-nuclei were calculated with the Wilcoxon rank sum test. Genes were considered differentially expressed with a Bonferroni-corrected p-value <0.05.

### Computation of neighboring CpG methylation states (see flow chart below)

Methylated cytosine counts per reads were computed using an in-house pipeline (bam_to_meCount.py) provided in the supplementary material. The output for bam_to_meCount.py script included the following columns in a table: Column1: read.names, Column 2: The number of Cs methylated in CpG context, Column 3: The number of Cs unmethylated in CpG context, Column 4: The number of Cs methylated in non-CpG context, Column 5: The number of Cs unmethylated in non-CpG context, Column 6: number of bases that are not Cs, Column 7: read.length. From this table, the proportion of methylated CpGs was computed. To get the coordinates per read, bam files were converted to bed files using bamtobed function. The converted bed files were concatenated with the bam_to_meCount.py table output. To get the reads that overlap the IMRs, we used intersectBed function. Then reads with a minimum of 2 CpGs were extracted for plotting. To normalize the reads per IMR, the proportion of reads per IMR per methylation state was computed using the prop.table function (data.table package, R). The distribution of read proportion per IMR methylation state was plotted using boxplot function (R).

### Ingenuity Gene Ontology Analysis

Lists of genes with IMRs and DM CpGs between FOS+ and FOS-cells, as well as with genes with DE were generated and used as input for Ingenuity Pathway Analysis (IPA). Top molecular/cellular functional categories, and functions within categories, were listed according to their corrected B-H value.

### Statistical Analysis

One-way, two-way, or repeated-measures ANOVAs with Sidak posthoc and paired and unpaired *t* tests were used. See statistics for DNA methylation and gene expression in corresponding sections.

## Supplementary Figures

**Supplementary Figure 1.**
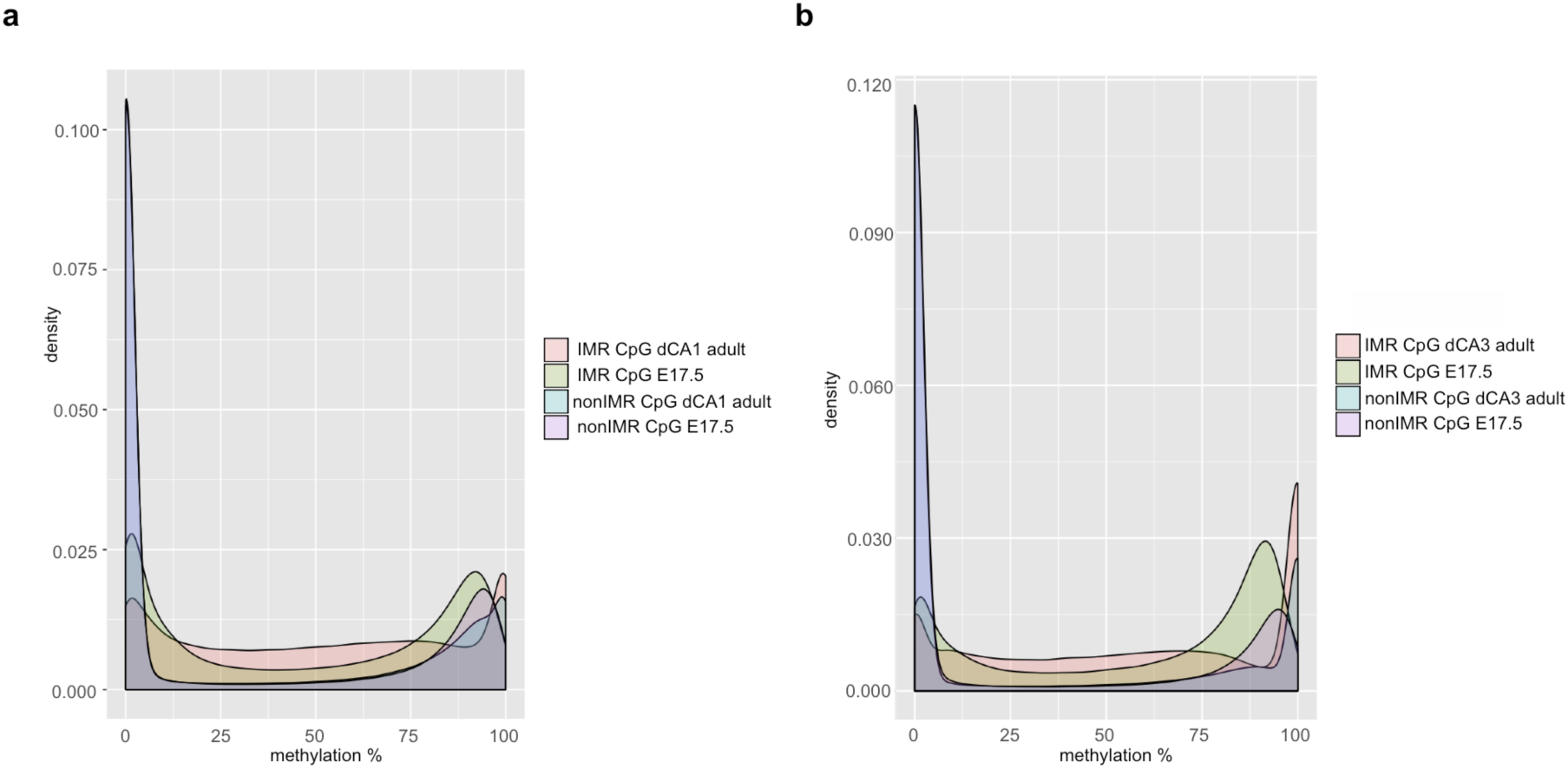
In addition to DGCs, IMRs are present in CA neurons in the hippocampus. **a, b.** Methylation distribution of IMR and non-IMR CpGs of CA1 and CA3 pyramidal cells at developmental time points E17.5 and adulthood in males.

**Supplementary Figure 2.**
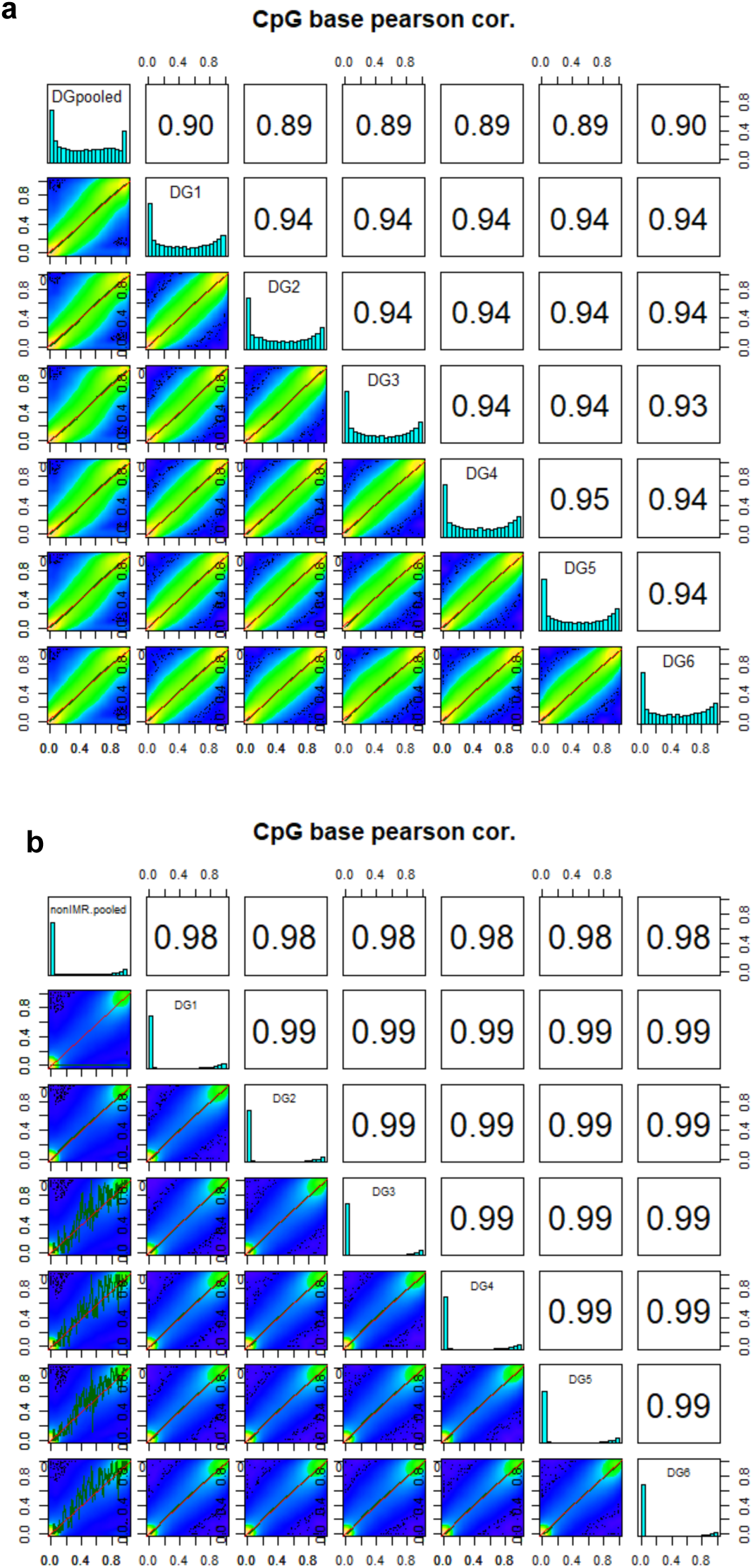
Correlation plots of methylation distribution at IMRs in DGCs of six individual animals. **a.** Single animal methylation distribution at IMR CpGs highly correlate with that of IMR CpGs from pooled samples, suggesting intermediacy in methylation at IMRs is not due to inter-animal differences in pooled samples. Individual profiles are also highly similar to each other. **b.** CpGs outside of IMRs exhibit the expected bimodal distribution in methylation.

**Supplementary Figure 3.**
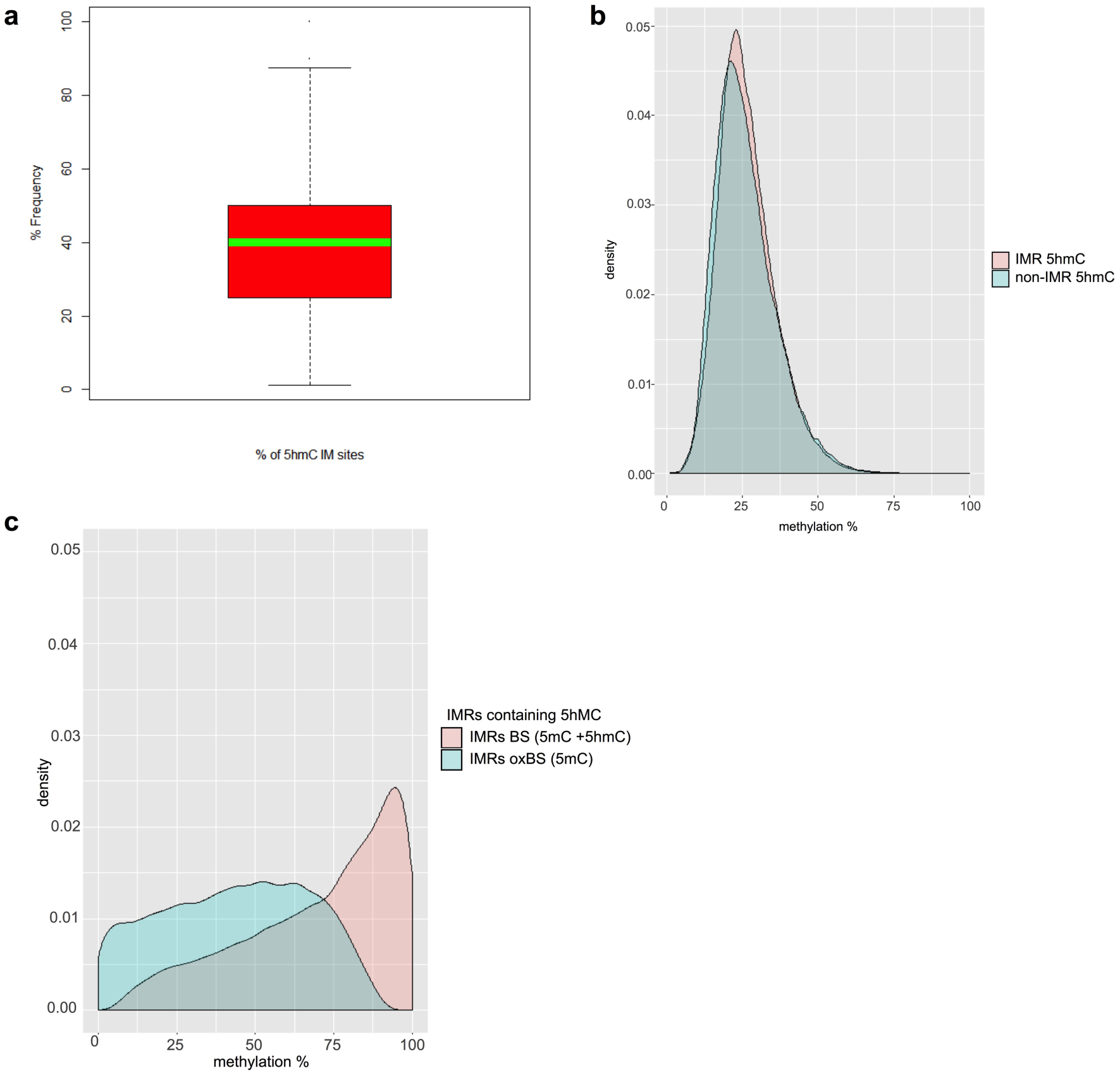
Characteristics of IMRs. **a.** OxBS sequencing reveals that only a fraction of IMRs contain 5hmC, with an average of 42.15% hydroxymethylation across IM sites. Boxplot of % frequency of 5hmC at IMRs with 5hmC, green line is median. **b.** Distribution of hydroxymethylated CpGs in IMR and non-IMR regions are comparable. **c.** Density plot showing methylation distribution of 5hmC containing IMRs. BS: 5hmC+5mC (pink) and oxBS: 5mC (blue). Methylation distribution of hydroxymethylated IMRs is IM after removal of 5hmC sites.

**Supplementary Figure 4.**
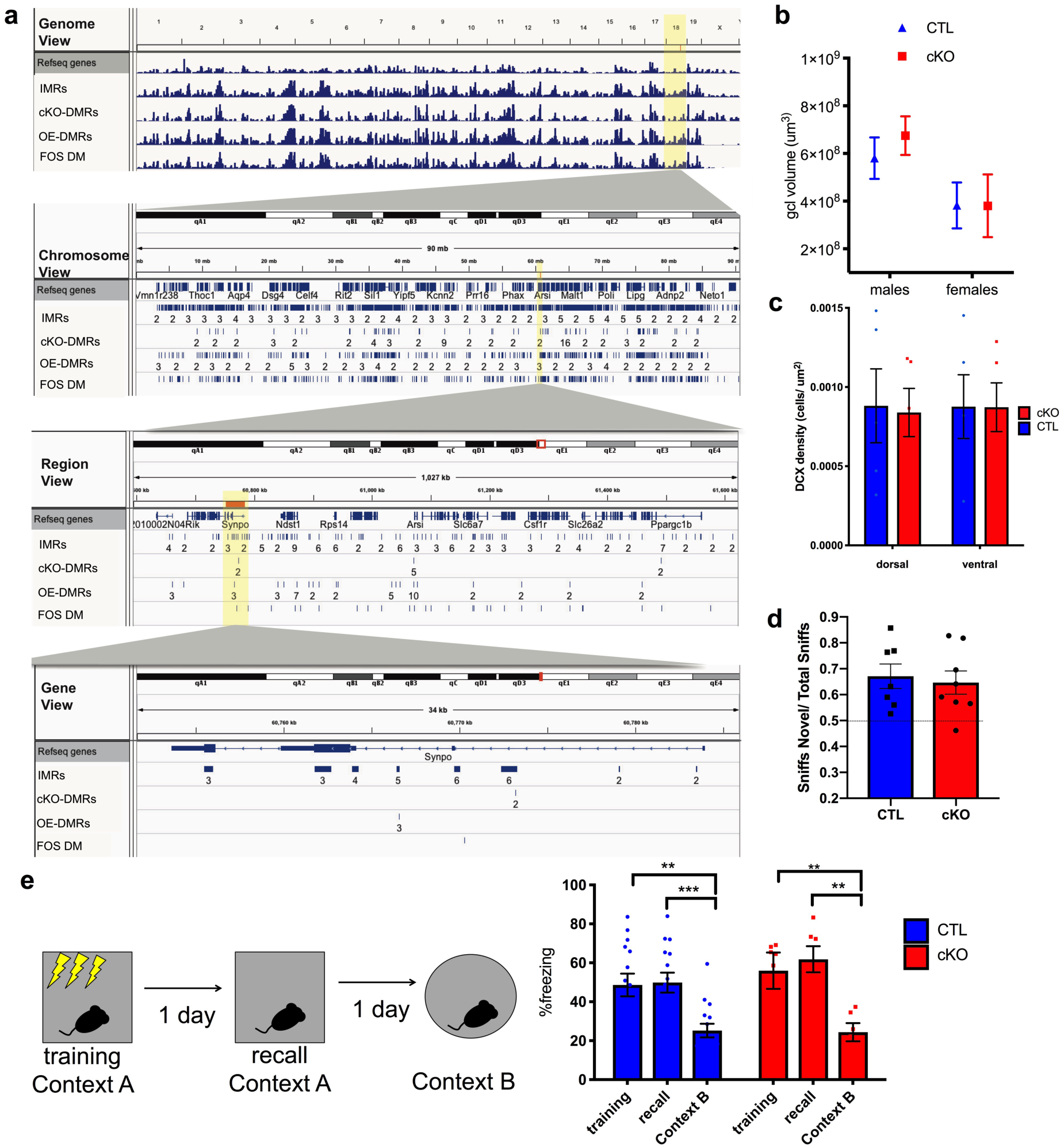
Conditional inactivation of *Dnmt3a* during development hypomethylates IMR CpGs. **a.** IGV Browser view of custom tracks under the mm9 gene track demonstrates the overlap between developmental IMRs and regions hypomethylated in cKO cells (cKO-DMRs), as well as regions hypermethylated in DNMT3A overexpressing cells (OE-DMRs) and sites differentially methylated in FOS^+^, relative to FOS^-^ cells (FOS DM). Overlap is apparent at the whole autosomal genome level (top), at chromosome level (Chr 18, middle), and at gene level (*Synpo*). **b,c.** Inactivation of *Dnmt3a* does not alter granule layer volume and adult neurogenesis in the DG. Granule cell layer volume in cKO and control (CTL) males and females. t-test: males, t(6)=0.8915, p=0.406992, n=4,4; females, t(8)=0.01098, p=0.991509, n=4,6). **c.** Comparable density of DCX positive young cells in the granule cell layer of adult cKO and control male mice. Multiple t-tests: dorsal t(8)=0.1535, p=0.881, n=5,5 mice, ventral t(8)=0.01532, p=0.988, n=5,5 mice). **d.** cKO mice perform similar to controls in novel object preference task. Ratio of sniffs to novel to total sniffs (novel object + habituated object). Both, control mice and cKO mice show novel object preference above chance (One sample t-test: control, t(1)=3.611, *p=0.0112); cKO, t(1)=3.243, *p=0.0142) There was no difference in control and cKO performance (Welch-corrected t=0.3756, p=0.7133, n=6,8). **e.** Control and cKO males froze at similar levels in context only fear conditioning training and recall, and both were able to discriminate between training and new context B (Two-way ANOVA genotype x day, F_(2,38)=_1.170, P=0.314; day, F_(2,38)_=32.30, P=<0.0001, N=15; 6). Significant difference between recall vs. context B of Controls is also displayed: adjusted ***p<0.0001. recall vs. context B of cKO: **p=0.0025; training vs. context B in Controls: adjusted *p=0.0028; training vs. context B in cKO: adjusted *p=0.0106. Data are mean ± s.e.m.

**Supplementary Figure 5.**
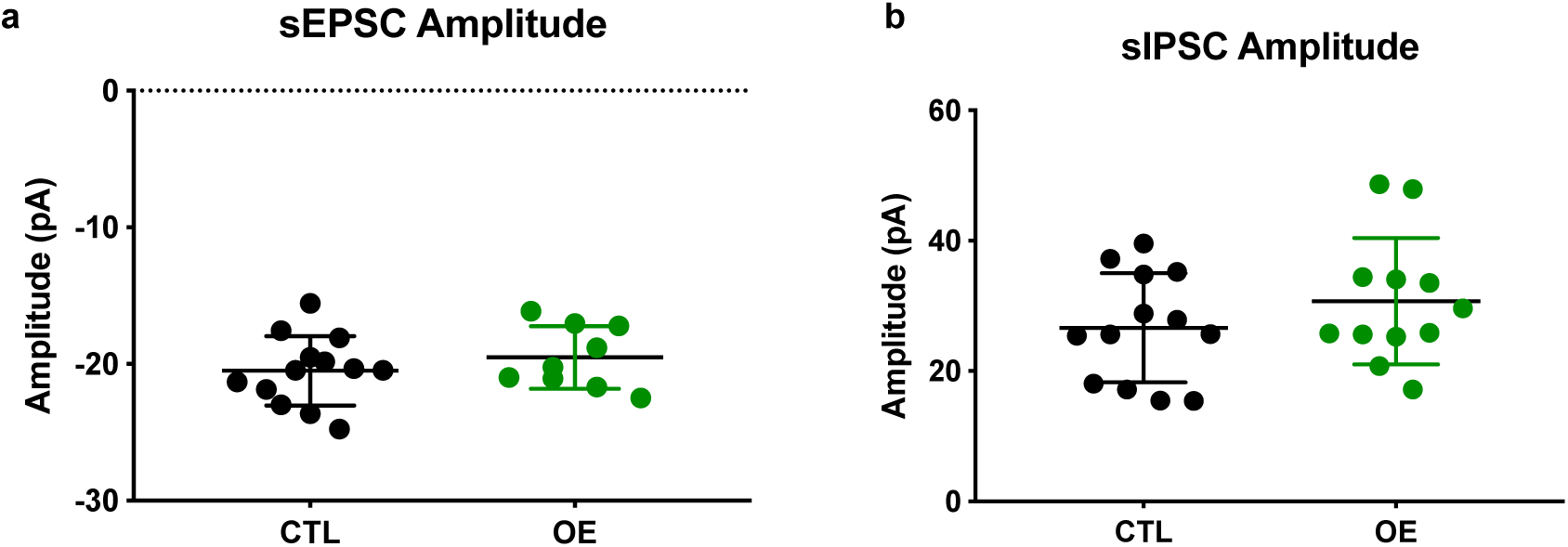
Synaptic transmission is not altered in DGCs overexpressing DNMT3A. **a, b.** Amplitude of spontaneous EPSCs and IPSCs were not significantly different in OE and control DGCs, indicating no detectable change in excitatory or inhibitory synaptic transmission. (N=control:14 cells/7 mice, OE:17 cells/6 mice).

