## Supplementary Table 2 for "Epigenetically bistable regions across neuron-specific genes govern neuron eligibility to a coding ensemble in the hippocampus"

**Supplementary Table 2. Links to bed files of IMRs.** Please install IGV on your local computer. Open IGV by clicking on “File”, then choose “Load from URL”. Load the bed file by copying the link of interest and pasting it into IGV.

| # | Region/sex | Link |
| --- | --- | --- |
| 1 | dDGC IMRs males | <https://abc.med.cornell.edu/bambam_athena/stream/track/1003.bed?access_token=477dd8d4a5c6bfc248ef41daefeb96ae> |
| 2 | dCA1 IMRs males | https://abc.med.cornell.edu/bambam_athena/stream/track/878.bed?access_token=22874f70f0d3d877971d577e45737d3f |
| 3 | dCA3 IMRs males | https://abc.med.cornell.edu/bambam_athena/stream/track/870.bed?access_token=f5adb2bd7d393052533e58ea81db3c9c |
| 4 | OE-DMRs dDGC males | https://abc.med.cornell.edu/bambam_athena/stream/track/876.bed?access_token=ad5b959c1140aeada9137fb38f191781 |
| 5 | cKO-DMRs dDGC males | https://abc.med.cornell.edu/bambam_athena/stream/track/859.bed?access_token=d5e05f720a69f3564f7d95a419f00b5d |
| 6 | DM CpGs FOS+/FOS- | <https://abc.med.cornell.edu/bambam_athena/stream/track/1016.bed?access_token=29ebb9a7fde1e26d168d653e385b1081> |
